# A human antibody reveals a conserved site on beta-coronavirus spike proteins and confers protection against SARS-CoV-2 infection

**DOI:** 10.1101/2021.03.30.437769

**Authors:** Panpan Zhou, Meng Yuan, Ge Song, Nathan Beutler, Namir Shaabani, Deli Huang, Wan-ting He, Xueyong Zhu, Sean Callaghan, Peter Yong, Fabio Anzanello, Linghang Peng, James Ricketts, Mara Parren, Elijah Garcia, Stephen A. Rawlings, Davey M. Smith, David Nemazee, John R. Teijaro, Thomas F. Rogers, Ian A. Wilson, Dennis R. Burton, Raiees Andrabi

## Abstract

Broadly neutralizing antibodies (bnAbs) to coronaviruses (CoVs) are valuable in their own right as prophylactic and therapeutic reagents to treat diverse CoVs and, importantly, as templates for rational pan-CoV vaccine design. We recently described a bnAb, CC40.8, from a coronavirus disease 2019 (COVID-19)-convalescent donor that exhibits broad reactivity with human beta-coronaviruses (β-CoVs). Here, we showed that CC40.8 targets the conserved S2 stem-helix region of the coronavirus spike fusion machinery. We determined a crystal structure of CC40.8 Fab with a SARS-CoV-2 S2 stem-peptide at 1.6 Å resolution and found that the peptide adopted a mainly helical structure. Conserved residues in β-CoVs interacted with CC40.8 antibody, thereby providing a molecular basis for its broad reactivity. CC40.8 exhibited in vivo protective efficacy against SARS-CoV-2 challenge in two animal models. In both models, CC40.8-treated animals exhibited less weight loss and reduced lung viral titers compared to controls. Furthermore, we noted CC40.8-like bnAbs are relatively rare in human COVID-19 infection and therefore their elicitation may require rational structure-based vaccine design strategies. Overall, our study describes a target on β-CoV spike proteins for protective antibodies that may facilitate the development of pan-β-CoV vaccines.

**SUMMARY:** A human mAb isolated from a COVID-19 donor defines a protective cross-neutralizing epitope for pan-β-CoV vaccine design strategies

## Introduction

Severe acute respiratory syndrome coronavirus 2 (SARS-CoV-2) has led to the current global pandemic (*1–3*). SARS-CoV-2 is a virus that belongs to the coronaviridae family of which six members have previously crossed into humans from animal reservoirs and established widespread infections (*4, 5*). These include four endemic human coronaviruses (HCoVs) (HCoV-229E, HCoV-HKU1, HCoV-OC43, HCoV-NL63) responsible for non-severe, seasonal infections (*4*) as well as SARS-CoV-1 and MERS-CoV (Middle East Respiratory Syndrome CoV) that are associated with high morbidity and mortality in humans (*6, 7*). Among the seven HCoVs, SARS-CoV-2 closely resembles SARS-CoV-1 and, to lesser degree, MERS-CoV. Together with HCoV-HKU1 and HCoV-OC43, these viruses belong to the *β-coronavirus* genus (*4, 5*). SARS-CoV-2 is highly transmissible in humans and causes coronavirus disease-2019 (COVID-19), associated with severe respiratory failure leading to high morbidity and a reported mortality of about 0.7 to 2% of infected individuals worldwide (*2, 8, 9*). There are considerable concerns that future coronavirus spillovers will trigger new pandemics (*10–15*).

Coronavirus pandemic preparedness may consider responses through establishment of techniques for rapid generation of specific reagents to counter the emerging coronavirus and control spread. An alternative is to seek to identify broadly neutralizing antibodies (bnAbs) to coronaviruses and use molecular information gleaned on their epitopes to rationally design pan-coronavirus vaccines (*16–18*). Pan-coronavirus vaccines and antibodies could be stockpiled ahead of the emergence of a new coronavirus and used to rapidly contain the virus. BnAbs and pan-coronavirus vaccines that target more conserved regions of the virus may also be more effective against antigenically variant viruses, such as have been described for the variants of concern in the COVID-19 pandemic (*19–22*).

All HCoVs possess a surface envelope spike glycoprotein that mediates interaction with host cell receptors and enables virus fusion (*4, 23*). SARS-CoV-2 (similar to SARS-CoV-1) utilizes the receptor binding domain (RBD) in the S1 subunit of the spike protein to engage human angiotensin converting enzyme 2 (hACE2) on host cells for cell entry and infection (*23–27*). The SARS-CoV-2 spike glycoprotein is the primary target of neutralizing antibodies (nAbs) (*28–31*). On the spike protein, the RBD is highly immunogenic and is recognized by the majority of nAbs (*28, 32–43*), and thus is a major focus of current nAb-based vaccine design efforts (*28, 44, 45*). However, due to sequence diversity, cross-reactivity to the RBD region is limited, especially among emerging coronaviruses with pandemic potential (*10–13*). The most potent nAbs in humans during natural infection are typically raised to epitopes overlapping the ACE2 binding site (*32, 33, 42, 45, 46*). As the rapid spread of the SARS-CoV-2 virus continues, these epitopes are coming under strong immune selection pressure at the population level, leading to the selection of SARS-CoV-2 neutralization escape variants (*19-22, 47-49*). The relevant mutations may result in reduced effectiveness of vaccine-induced antibody responses in humans since such responses also tend to target RBD epitopes overlapping the ACE2 binding site, and because all currently approved vaccines are based on the wild-type virus. The most striking example that illustrates the capability of the RBD to mutate without majorly affecting the ability of the virus to engage host receptor is the variability of the RBD across the two families of HCoVs: SARS-CoV-2/1 (β-HCoVs) and HCoV-NL63 (α-HCoV) (*23–27, 50*). These HCoVs possess divergent RBDs, but all use the ACE2 receptor for viral entry suggesting that SARS-CoV-2, and potentially other emerging sarbecoviruses with human pandemic potential, can tolerate changes in this domain with limited fitness cost. Therefore, we believe that other sites on the spike protein should be explored as targets of bnAbs.

We recently isolated a SARS-CoV-1/2 cross-neutralizing antibody from a COVID-19 donor, CC40.8, that exhibits broad cross-reactivity with human β-CoVs (*51*). Here, we show that the CC40.8 bnAb targets an S2 stem-helix epitope, which is part of the coronavirus fusion machinery. We first identified a long 25-mer S2 peptide from HCoV-HKU1 that bound CC40.8 with high affinity and then determined the crystal structure of CC40.8 with the SARS-CoV-2 S2 peptide. The S2 stem peptide adopts a largely helical structure that is embedded in a groove between the heavy and light chain complementarity determining regions (CDRs) of the antibody. Key epitope contact residues were further validated, by alanine scanning, to be important for peptide binding and for virus neutralization. These contact residues are largely conserved between β-CoVs, consistent with the cross reactivity of CC40.8. In SARS-CoV-2 challenge models, CC40.8 showed in vivo protective efficacy by reducing weight loss and lung tissue viral titers. Although two recent studies have described S2-stem nAbs isolated from mice and mice transgenic for human Ig (*52, 53*), CC40.8 represents a human HCoV S2-stem directed bnAb isolated from natural infection (*51*) and may facilitate development of antibody-based interventions and prophylactic pan-sarbecovirus and pan-β-coronavirus vaccine strategies.

## Results

### CC40.8 binds a conserved peptide from the S2 region of β-coronaviruses

We recently isolated a bnAb, CC40.8, from a 62-year-old SARS-CoV-2 convalescent donor from peripheral blood mononuclear cell (PBMC) samples collected 32 days post-infection (*51*). CC40.8 bnAb neutralizes SARS-CoV-1 and SARS-CoV-2 and exhibits broad reactivity against β-coronaviruses, notably the endemic coronavirus HCoV-HKU1 (Fig. 1A and B) (*51*). Here, we observed that CC40.8 bnAb can effectively neutralize clade 1b and clade 1a ACE2 receptor-utilizing sarbecoviruses (Fig. 1A, fig. S1A). In addition, the CC40.8 bnAb was consistently effective against the current SARS-CoV-2 variants of concern (VOCs) (Fig. 1A, fig. S1A). The effectiveness of CC40.8 bnAb with SARS-CoV-2 VOCs is consistent with a lack of mutations in the S2 stem helix region in the current VOCs (*21*). To assess the cell-cell inhibition ability of CC40.8 bnAb, we conducted experiments in HeLa cells expressing SARS-CoV-2 spike protein or hACE2 receptor. We observed that CC40.8 bnAb can prevent cell-cell fusion of HeLa cells expressing SARS-CoV-2 spike protein with HeLa cells expressing the hACE2 receptor (fig. S2).

**Figure 1.**
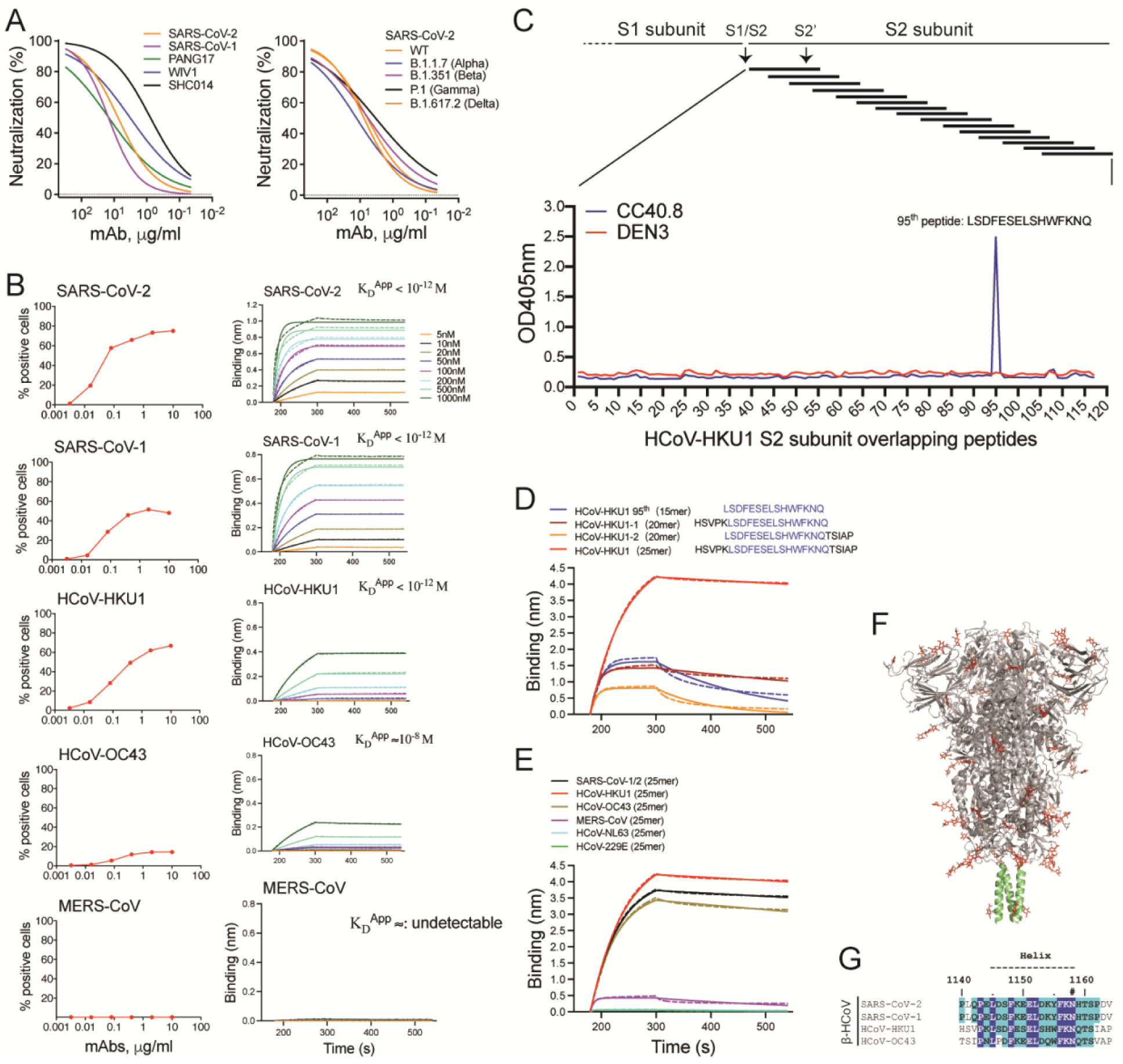
Identification of the CC40.8 bnAb epitope on the coronavirus spike protein by epitope mapping. **(A)** Neutralization of clade 1b (SARS-CoV-2 and Pang17) and clade 1a (SARS-CoV-1, WIV1 and SHC014) ACE2-utilizing sarbecoviruses by CC40.8 mAb isolated from a COVID-19 donor (left) is shown. CC40.8 neutralizing activity against SARS-CoV-2 (WT-Wuhan) and SARS-CoV-2 variants of concern [B.1.1.7 (alpha), B.1.351 (beta), P.1 (gamma) and B.1.617.2 (delta)] is shown on the right. **(B)** Left: Cellular ELISA (CELISA) data show binding of CC40.8 mAb with β-HCoV spikes expressed on 293T cells. Binding to HCoV spikes is recorded as % positive cells using flow cytometry. CC40.8 mAb shows cross-reactive binding with 4 out of 5 human β-HCoV spikes. Right: BioLayer Interferometry (BLI) binding of CC40.8 mAb with human β-HCoV soluble spike proteins. Apparent binding constants (KD) for each Ab-antigen interaction are indicated. KD^App^ <10^-12^M indicates that no off-rate could be measured. The raw experimental curves are shown as dash lines, while the solid lines are the fits. **(C)** Epitope mapping of CC40.8 with HCoV-HKU1 S2 subunit overlapping peptides is shown. A series of HCoV-HKU1 S2 (GenBank: AAT98580.1) overlapping biotinylated peptides (15-residues long with a 10-residue overlap) were tested for binding to CC40.8 mAb by ELISA. OD405, optical density at 405nm. CC40.8 showed binding to the 95^th^ 15-mer peptide corresponding to the HCoV-HKU1 S2 stem-helix region (residue position range: 1231-1245). An antibody to dengue virus, DEN3, was used as a control. **(D)** BLI data are shown for CC40.8 binding to the HCoV-HKU1 95^th^ 15-mer stem peptide (blue) and HCoV-HKU1 stem peptide variants with 5 additional residues either at the N-(20-mer: brick red) or C-(20-mer: orange) terminus or added at both termini (25-mer: red). CC40.8 showed strongest binding to the 25-residue stem peptide corresponding to HCoV-HKU1 S2 residues 1226-1250. The kinetic curves are fit with a 1:1 binding mode. **(E)** BLI data are shown for CC40.8 binding to 25-mer stem peptides derived from different HCoV spikes. CC40.8 showed binding to the β- but not to the α-HCoV S2 stem peptides. The HCoV-HKU1 S2 residues 1226-1250 correspond to residues 1140-1164 on SARS-CoV-2 spike. The kinetic curves are fit with a 1:1 binding mode. **(F)** A SARS-CoV-2 spike protein cartoon depicts the S2-stem epitope region in green at the base of the prefusion spike ectodomain. **(G)** Sequence conservation of the CC40.8 stem-helix epitope is shown for SARS-CoV-1/2, HCoV-HKU1 and HCoV-OC43 human β-CoV spike proteins. Conserved identical residues are highlighted with blue boxes, and similar residues are in cyan boxes [amino acids scored greater than or equal to 0 in the BLOSUM62 alignment score matrix (*92*) were counted as similar here]. An N-linked glycosylation site is indicated with a “#” symbol.

Using negative-stain electron microscopy (ns-EM), we previously showed that the CC40.8 antibody targets the base of the S2 subunit on HCoV spike proteins, but epitope flexibility precluded determination of a high-resolution structure (*51*). Here, we pursued epitope identification, first by peptide mapping. Using HCoV-HKU1 S2 subunit overlapping biotinylated peptides (15-residue long with a 10-residue overlap) for binding to CC40.8, we identified that the stem-helix region in the S2 fusion domain contains the epitope (Fig. 1C, fig. S3). Then, through screening with peptides of various lengths that include the epitope, we identified a 25-residue peptide that showed the strongest binding by biolayer interferometry (BLI, Fig. 1D). The peptide corresponds to residues 1226-1250 from the HCoV-HKU1 S2 sequence.

Next, we tested BLI binding of CC40.8 bnAb with peptides encompassing similar S2-domain regions of other HCoVs. We observed that the antibody binds to the β- but not to the α-HCoV S2-domain peptides (Fig. 1D). This pattern is consistent with the differential binding of CC40.8 bnAb to different families of HCoV spike proteins (Fig. 1B) (*51*). Sequence alignment of the S2 stem-helix domain region showed strong conservation between SARS-CoV-1 and SARS-CoV-2 with more modest conservation across the seasonal β-CoVs, consistent with cross-reactive binding patterns (Fig. 1E to G).

To determine whether CC40.8 bnAb affinity maturation was important for cross-reactive binding or neutralization, we generated an inferred germline (iGL) version of CC40.8 with corresponding antibody V-D-J germline genes (fig. S1B), as described previously (*54, 55*). Although the CC40.8 bnAb iGL Ab version retained binding to spike proteins and the stem-helix peptides of β-CoVs, the neutralizing activity was lost against sarbecoviruses (fig. S1, C to E), suggesting that affinity maturation is critical for neutralization, but cross-reactive breadth is germline-encoded. Interestingly, the CC40.8 iGL bnAb showed binding to MERS-CoV spike protein that CC40.8 bnAb fails to bind and exhibited some weak polyreactivity (fig. S1, fig. S4), suggesting that naive B cells targeting this epitope may begin with a broader reactivity to CoV spike proteins.

### The epitope of CC40.8 bnAb was defined by the crystal structure of a peptide-antibody complex

To investigate the molecular nature of the CC40.8 bnAb epitope, we determined the 1.6 Å resolution crystal structure of the antibody Fab fragment with the SARS-CoV-2 25-mer S2 peptide (Fig. 2A, table S1). The peptide adopts a largely helical structure that traverses a wide hydrophobic groove formed between the heavy and light chains of the Fab (fig. S5). The buried surface area on the peptide is about 1150 Å^2^ (669 Å^2^ conferred by the heavy chain and 488 Å^2^ by the light chain) and is largely contributed by hydrophobic residue interactions at the paratope-epitope interface, although some hydrogen bonds and salt bridges are contributed by CDRHs1 to 3, FRH1, and CDRLs1 to 3 (Fig. 2A to C, fig. S6). Two peptide stretches of ^1142^QPELD^1146^ and ^1151^ELDKYF^1156^ and several nearby residues, F^1148^ N^1158^, H^1159^, form the epitope of the bnAb (Fig. 2B and C). Notably, hydrophobic residues in ^1151^ELDKYF^1156^ of the stem region, as well as two upstream residues, L^1145^ and F^1148^, form the core of the epitope that interacts with a hydrophobic groove in the antibody lined by heavy chain residues (V33, Y35, W47, Y56, Y58, M96 and V101) and light chain residues (Y32, Y34, L46 and Y49) (Fig. 2C and fig. S6). Antibody germline and mutated residues both contribute to epitope recognition (fig. S6). Consistent with our findings, two recent independent studies have shown that heterologous CoV spike protein immunizations in mice or mice transgenic for the human Ig locus can induce cross-reactive serum neutralizing antibodies that target the conserved S2 spike epitope similar to the stem-helix epitope identified in our study, and some isolated mAbs also show broad reactivity to coronavirus spike proteins (*52, 53*).

**Figure 2.**
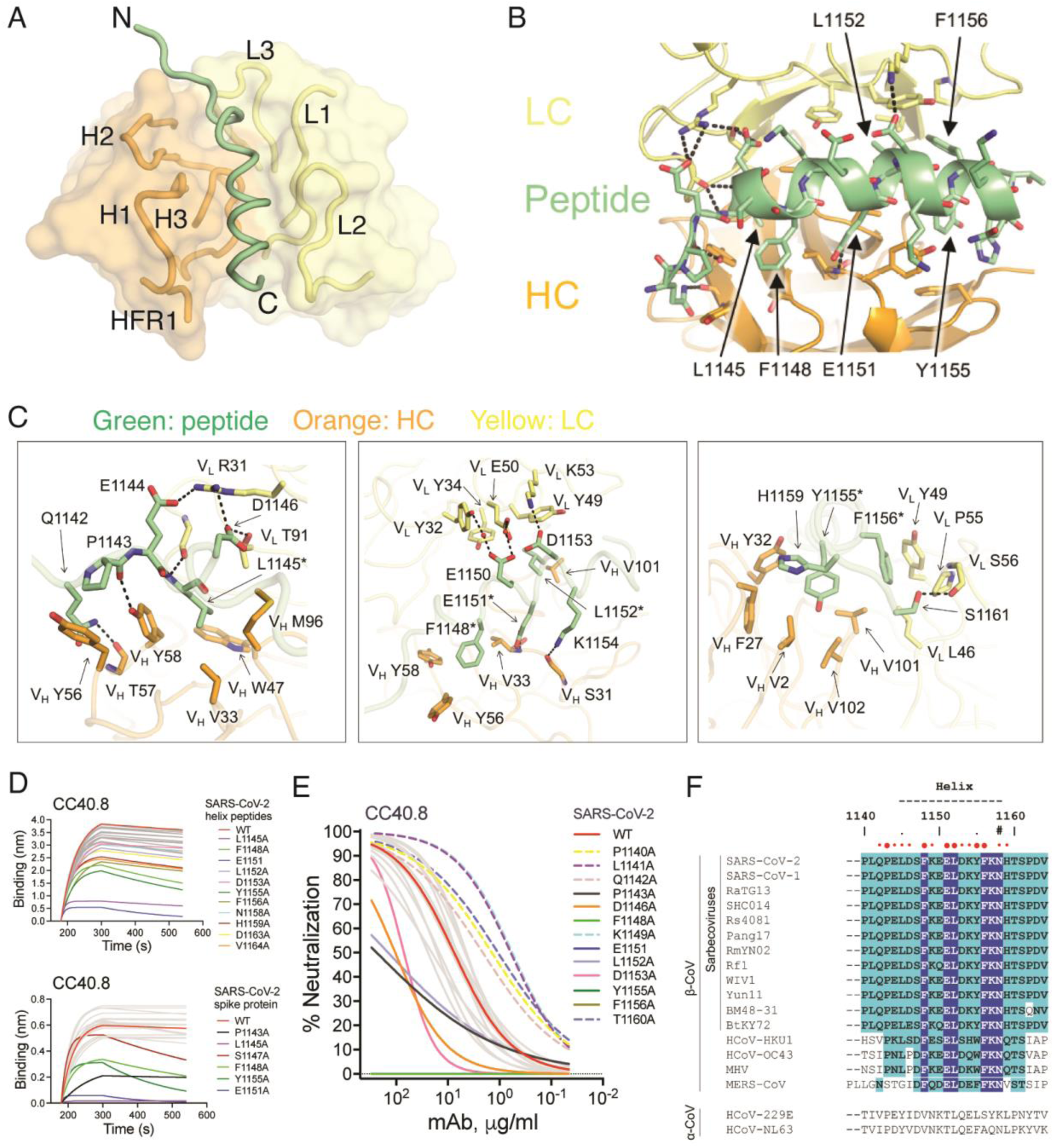
Crystal structure of CC40.8 antibody in complex with the SARS-CoV-2 stem peptide, and S2 stem bnAb epitope residues and conservation across CoVs. **(A)** An overall view of the CC40.8-peptide complex structure is shown at 1.6 Å resolution. Heavy and light chains of CC40.8 are shown in orange and yellow semi-transparent surfaces, respectively, with the heavy (H) and light (L) chain complementary determining regions (CDRs) shown as tubes. The SARS-CoV-2 stem-helix peptide is shown as a green tube for the peptide backbone. **(B)** An overview of the CC40.8 antibody and S2 stem-peptide interaction is shown. Heavy (H) and light (L) chains of CC40.8 are shown in orange and yellow, respectively, whereas the SARS-CoV-2 stem peptide is in green. Hydrogen bonds and salt bridges are represented by black dashed lines. **(C)** Details of the interactions between CC40.8 and the SARS-CoV-2 stem peptide are shown. Residues conserved in SARS-CoV-1, SARS-CoV-2, and other sarbecoviruses as well as seasonal β-CoVs HCoV-HKU-1, and HCoV-OC43 are labeled with asterisks (*). **(D)** BLI data are shown for binding of CC40.8 bnAb to SARS-CoV-2 stem-helix peptide (top) and soluble spike protein alanine mutants (bottom) spanning the whole epitope. The stem peptide or spike protein mutants that substantially affect CC40.8 bnAb binding are shown in assorted colors in comparison to wild-type (WT, red). **(E)** Neutralization of SARS-CoV-2 and the stem-helix alanine mutants spanning the whole epitope by CC40.8 is shown. The WT virus is shown in red and virus mutants that substantially affect CC40.8 bnAb neutralization are shown in assorted colors. The bold and dashed color curves indicate substitutions that, respectively, led to a decrease or an increase in the IC50 neutralization titers compared to WT virus. **(F)** Sequence conservation is shown for the CC40.8 stem-helix epitope on SARS-CoV-1/2, sarbecoviruses infecting other animal species, human β-CoVs and mouse hepatitis virus (MHV). The stem region forming the helix is indicated by black dashes and residues involved in interaction with CC40.8 antibody are indicated by red dots (cutoff distance = 4 Å). Larger dots indicate residues that are essential for CC40.8 interaction as defined by alanine scanning mutagenesis where mutation decreased neutralization IC50 by at least 10-fold or a complete knock-out (details are shown in fig. S7). Conserved identical residues are highlighted with blue boxes, and similar residues are in cyan boxes [amino acids scored greater than or equal to 0 in the BLOSUM62 alignment score matrix (*92*) were counted as similar here]. An N-linked glycosylation site is indicated with a “#” symbol. The region that presents a helical secondary structure in the CC40.8/peptide structure is indicated on top of the panel.

**Figure 3.**
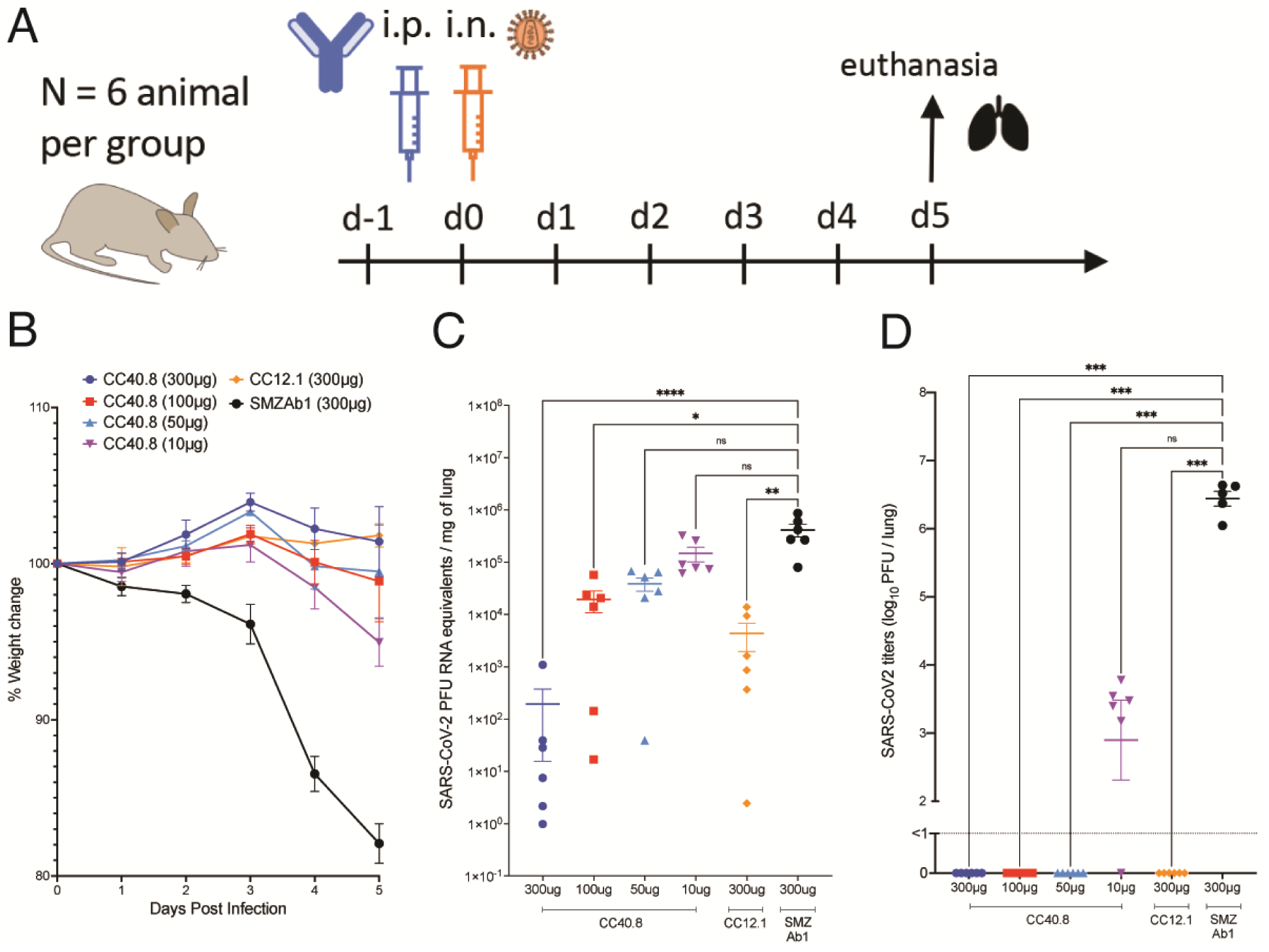
CC40.8 reduces weight loss, lung viral load, and viral replication following SARS-CoV-2 challenge in the hACE2 mouse model. **(A)** CC40.8 was administered intraperitonially (i.p.) at four different doses (300 μg, 100 μg, 50 μg, and 10 μg) per animal into hACE2 receptor-expressing mice (6 animals per group). Control animals received CC12.1 RBD nAb (300 μg per animal) or a Zika-specific mAb, SMZAb1 (300 μg per animal). Each group of animals was challenged intranasally (i.n.) 12 hours after antibody infusion with 2 × 10^4^ PFU of SARS-CoV-2 (USA-WA1/2020). Animal weight was monitored daily as an indicator of disease progression and lung tissue was collected on day 5 for viral load and viral burden assessment. **(B)** Percent weight change in CC40.8 or control antibody-treated animals after SARS-CoV-2 challenge is shown. Percent weight change was calculated from day 0 for all animals. Data are presented as mean ± SEM. **(C)** SARS-CoV-2 viral RNA loads based on the qPCR analysis of lung tissue at day 5 after infection are shown. Data are presented as mean ± SEM. **(D)** SARS-CoV-2 infectious virus titers (plaque-forming unit (PFU)) are shown as determined by plaque assay from lung tissue at day 5 after infection. Data are presented as mean ± SEM. Statistical comparisons between groups were performed using a Kruskal-Wallis non-parametric test followed by Dunnett’s multiple comparisons. (*p <0.05, **p <0.01, ***p <0.001; ****p < 0.0001; ns, p >0.05).

The residues important for CC40.8 interaction with virus were also investigated by alanine scanning mutagenesis of SARS-CoV-2 and HCoV-HKU1 peptides and spike protein by antibody binding and by neutralization of SARS-CoV-2 spike protein mutants (Fig. 2D and E, fig. S7). The contact residues determined by crystallography were also found to be important for peptide binding and neutralization with the S2 helical residues, L^1145^, E^1151^, F^1148^ and Y^1155^, being the most critical (fig. S7). We noted some differences in CC40.8 dependence on S2 residue substitutions for virus neutralization and spike protein binding, which may reflect differences in conformation or glycosylation between recombinant and native membrane-associated spike protein (*56, 57*). The conservation of those residues identified by crystallography and alanine scanning as most critical for interaction of CC40.8 with virus is high (Fig. 2F) across human β-coronaviruses and related sarbecoviruses that infect various animal species, thus providing a structural basis for broad cross-reactivity of the antibody.

The CC40.8 epitope region houses an N-linked glycan (N^1158^) that is highly conserved across coronaviruses and may restrict access to this bnAb epitope. To investigate, we substituted the T^1160^ residue on SARS-CoV-2 virus spike protein with an alanine residue to eliminate the AsnHisThr (NHT) N-linked glycan attachment site. A modest increase in neutralization sensitivity of the T1160A variant relative to wild-type virus was observed (Fig. 2E, fig. S7), suggesting that any steric obstruction of the epitope by the N1158 glycan is limited.

The CC40.8 bnAb epitope appears to be only partially exposed on the pre-fusion HCoV spike protein (fig. S8). Previously, a SARS-CoV-2 S2 stem-targeting neutralizing antibody, B6, was isolated from a mouse immunized with spike protein (*52*) . Here we compared the structures of antibodies CC40.8 and B6 (fig. S9, A to E). Both antibodies target a similar epitope on the SARS-CoV-2 spike protein, the conserved S2 stem helix region, but with different angles of approach; a longer peptide was visualized as the epitope for CC40.8. The post-fusion spike protein requires a large conformational change in the S2 stem region, and the superimposed CC40.8 (fig. S8C) and B6 (*52*) would clash with the post-fusion conformation. On the other hand, if bound to a spike protein in the pre-fusion state, both antibodies would clash with the adjacent protomer (fig. S9, D and E), suggesting a possible neutralization mechanism where the antibodies may induce disruption of the S2 stem 3-helix bundle, and bind to an intermediate state of the spike protein (fig. S8C). This hypothesis is further supported by comparing the binding of CC40.8 to S2P and HexaPro (S6P). The two proline mutations (S2P) were introduced to stabilize the SARS-CoV-2 S trimer in its pre-fusion state (*26, 58*), whereas additional proline substitutions to the HexaPro or S6P construct further stabilized the SARS-CoV-2 spike trimer (*59*). Here we show that the S6P-stabilized version exhibited much weaker binding to CC40.8 compared to S2P (fig. S9F), further suggesting that destabilization or partial disruption of the pre-fusion S trimer is a possible explanation for neutralization by S2 stem-targeting antibodies, such as CC40.8 or B6. Our previous EM study of the complex of HCoV-HKU1 S (S2P) and CC40.8 Fab [Fig. 5D in ref (*51*)] showed high flexibility of the epitope and multiple antibody approach angles, which also suggested disruption of the 3-helix bundle and induction of flexibility in the S2 stem region.

### CC40.8 antibody protects against weight loss and reduces viral burden in SARS-CoV-2 challenge in vivo

To determine the in vivo efficacy of CC40.8, we conducted passive antibody transfer followed by SARS-CoV-2 challenge in human ACE2 (hACE2) mice and in Syrian hamsters. CC40.8 mAb at 4 different doses (300μg, 100μg, 50μg and 10μg per animal) was intra-peritoneally (i.p.) administered into groups (6 animals per group) of hACE2 mice (Fig. 3A) (*60*). An RBD nAb (CC12.1; 300 μg/animal) positive control and a Zika-specific antibody (SMZAb1; 300 μg/animal) negative control were administered i.p. into control animal groups. All CC40.8- and control mAb-treated animals were challenged with SARS-CoV-2 (USA-WA1/2020) by intranasal (i.n.) administration of a virus dose of 2 × 10^4^ plaque forming units (PFU), 12 hours post-antibody infusion (Fig. 3A). The animals were weighed daily to monitor weight changes, as an indicator of disease due to infection and serum samples were collected to determine the transferred human antibody concentrations (Fig. S10). Animals were euthanized at day 5 and lung tissues were collected to determine the SARS-CoV-2 titers by quantitative polymerase chain reaction (qPCR) and by plaque assays. The CC40.8 bnAb-treated animals showed significantly reduced weight loss as compared to the SMZAb1-treated control group animals (P<0.0001, Fig. 3B, fig. S10), suggesting a protective role for CC40.8. Remarkably, the animals treated with the lowest dose of CC40.8 bnAb (10 μg/animal) also showed significant protection against weight loss (P = 0.0005, Fig. 3B, fig. S10). As expected, the positive control RBD nAb, CC12.1 significantly protected against weight loss (P<0.0001, Fig. 3B, fig. S10). Consistent with these results, SARS-CoV-2 specific viral RNA copies and viral titers in day 5 lung tissues were significantly reduced in the CC40.8-treated animals compared to the SMZAb1 control group animals (P<0.0001, fig. 3C and D).

We also investigated the protective efficacy of CC40.8 mAb by intra-peritoneally (i.p.) administering into a group of 5 Syrian hamsters (at 2 mg per animal) and subsequently challenging with SARS-CoV-2 (USA-WA1/2020 dose of 1 × 10^6^ PFU) (fig. S11). SMZAb1 Zika mAb was used as a control. Consistent with hACE2 mouse experiments, the CC40.8 bnAb-treated animals showed substantially reduced weight loss and reduced SARS-CoV-2 titers in day 5 lung tissues demonstrating its protective role (fig. S11). Altogether, the findings reveal that CC40.8, despite relatively low in vitro neutralization potency, shows a substantial degree of protective efficacy against SARS-CoV-2 infection in vivo. Consistent with these results, a recently isolated S2 stem bnAb, S2P6, has also been shown to protect against SARS-CoV-2 challenge despite relatively low neutralization potency (*61*) . Furthermore, this phenomenon of a surprisingly high degree of protection afforded by antibodies directed to epitopes close to the spike protein membrane and part of the fusion machinery has been described earlier for HIV (*62*).

### The conserved stem-helix epitope defined by bnAb CC40.8 is infrequently targeted following SARS-CoV-2 infection

To investigate how frequently the CC40.8 epitope is targeted following SARS-CoV-2 infection, we tested the binding of serum samples from 60 COVID-19 convalescent donors to 25-mer peptides of HCoVs corresponding to the stem-helix bnAb epitope. We observed that 6 of 60 (10%) individuals exhibited some degree of cross-reactive binding with β-HCoV S2 stem peptides (Fig. 4A). We further tested the binding of cross-reactive serum samples with SARS-CoV-2 S2 stem peptide alanine scan variants spanning the CC40.8 epitope and observed the presence of CC40.8-like epitope-targeting antibodies (Fig. 4B). The binding of cross-reactive serum Abs revealed dependence on five common stem helix residues including a conserved hydrophobic core formed by F^1148^, L^1152^, Y^1155^ and F^1156^ (Fig. 4B). To determine the contribution of S2-stem directed antibodies in overall SARS-CoV-2 neutralization by serum Abs in cross-reactive COVID-19 donors, we conducted competition experiments with the SARS-CoV-2 S2 stem-helix peptide. Peptide competition showed no or minimal effects on the SARS-CoV-2 neutralization (Fig. 4C and D), suggesting that stem-helix targeting cross-reactive nAbs minimally contribute to the overall polyclonal serum neutralization in these COVID-19 convalescent donors.

**Figure 4.**
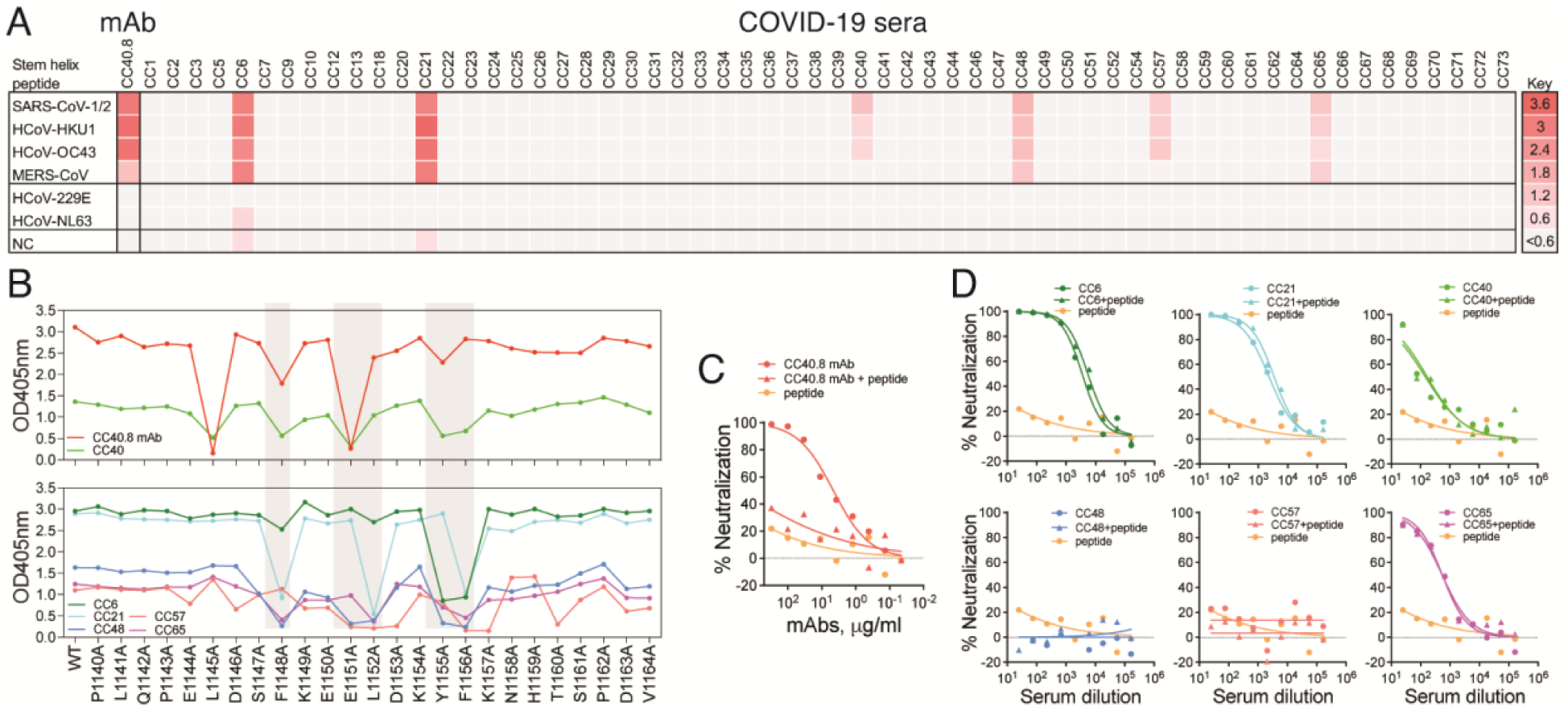
Frequency of CC40.8 S2 epitope-targeting serum antibodies in human COVID-19 donors. **(A)** The heatmap shows ELISA binding reactivity profiles of convalescent COVID-19 serum samples with 25-mer peptides corresponding to the CC40.8 bnAb S2 epitope on human β-(SARS-CoV-2, SARS-CoV-1, MERS-CoV, HCoV-HKU1, HCoV-OC43) and α-(HCoV-NL63 and HCoV-229E) coronaviruses. The extent of binding (represented as OD405 values) is color coded with red indicating strong reactivity. CC40.8 mAb was the positive control for the binding assay and PBS-BSA solution served as the negative control. Six out of 60 COVID-19 convalescent donors showed cross-reactive binding to various HCoV spike stem-helix peptides. **(B)** ELISA-based alanine scan epitope mapping is shown for convalescent COVID-19 serum samples from CC6, CC21, CC40, CC48, CC57 and CC65 donors with SARS-CoV-2 stem peptides (25mer). CC40 serum showed dependence on similar SARS-CoV-2 stem-helix residues as the CC40.8 mAb. SARS-CoV-2 stem-helix residue positions targeted (decrease in ELISA binding compared to WT stem peptide) by multiple cross-reactive COVID-19 serum samples are shown in gray. Five residues, F^1148^, E^1151^, L^1152^, Y^1155^ and F^1156^ were commonly targeted by the cross-reactive COVID-19 serum Abs. These residues form the stem-helix bnAb core epitope. **(C)** SARS-CoV-2 neutralization by CC40.8 in the presence of competing SARS-CoV-2 stem peptide is shown. Neutralization data are presented for SARS-CoV-2 by CC40.8 mAb, CC40.8 mAb pre-incubated with SARS-CoV-2 stem peptide (60 µg/ml) and stem peptide-only control. The SARS-CoV-2 stem peptide inhibits the neutralizing activity of CC40.8 mAb. **(D)** SARS-CoV-2 neutralization by cross-reactive COVID-19 serum samples was evaluated in the presence of competing SARS-CoV-2 stem peptide. Neutralization of SARS-CoV-2 by serum from COVID-19 convalescent donors, CC6, CC21, CC40, CC48, CC57, CC65, pre-incubated with SARS-CoV-2 stem peptide (60 μg/ml) and stem peptide-only controls was measured. The SARS-CoV-2 stem peptide had minimal effects on neutralization by these COVID-19 convalescent donor serum antibodies.

## Discussion

The development of effective pan-coronavirus vaccine strategies that can mitigate future outbreaks from new emerging coronaviruses is important (*16, 18*). Two major challenges are the identification of broadly neutralizing antibody (bnAb) targets on CoV spike proteins and the development of vaccine strategies that can reproducibly elicit durable and protective pan-CoV bnAbs. The approach of identifying conserved bnAb surface protein targets by isolating bnAbs from natural infection and utilizing their molecular information in structure-guided immunogen design has greatly contributed to the development of vaccine strategies against a range of complex pathogen surfaces (*63–70*).

The spike S1 subunit shows considerable variation on HCoVs, whereas the S2 subunit is relatively more conserved, especially across the β-HCoVs, and appears to be promising for developing pan-CoV bnAb vaccine strategies. Accordingly, we recently isolated a SARS-CoV-1/2 cross-neutralizing Ab, CC40.8, that exhibits broad reactivity with human β-CoVs (*51*). In this study, using epitope mapping and structural studies, we determined the spike epitope recognized by CC40.8. The epitope is located in the S2 stem-helix region, which is conserved across β-coronaviruses and may thus serve as a promising target for pan-β-coronavirus vaccine strategies. The epitope is highly enriched in hydrophobic residues as well as some charged residues. The bnAbs targeting this region may neutralize by sterically interfering with the fusion machinery (*52, 53*), suggesting a potential target for fusion inhibitors (*71–73*). CC40.8 bnAb represents a human bnAb directed to the HCoV S2 stem helix (*51*). Two more S2 stem human bnAbs, S2P6 and CV3-25, have also been reported recently (*61, 74*) that target a similar S2 stem epitope region. Knowledge from these nAbs will be important for developing broad vaccine strategies for β-coronaviruses.

We noted that cross-reactive antibodies directed to the CC40.8 S2 stem-helix epitope are much less frequently elicited in human coronavirus natural infections as compared to strain-specific neutralizing antibody responses (*28*). However, a few recent studies using more sensitive antibody detection assays have suggested a higher prevalence of polyclonal stem-helix region-directed antibodies in COVID-19 donors and their possible association with reduced disease severity (*75–77*). Regardless, the small subset of individuals in our sample cohort that do make cross-reactive Abs, seem to exhibit broad reactivity to human β-coronaviruses, which is promising for pan-β-coronavirus vaccine strategies. In principle, the paucity of these cross-reactive antibodies could be due to poor accessibility of the S2 stem-helix epitope on the native spike protein relative to other epitopes, low frequency of bnAb-encoding B cell precursors in humans, or complex secondary B cell maturation pathways. Low epitope accessibility is clearly a potential contributor to low immunogenicity. Low precursor frequency seems unlikely, at least for CC40.8-like antibodies given that this antibody uses a common VH gene segment (IGHV3-23) and CDRH3 length (10 amino acids) (*78*). Analysis of CC40.8 antibody variable regions by the Armadillo tool (*79*) revealed the presence of several improbable somatic mutations that are predicted to contribute to difficulty in elicitation of CC40.8-like antibodies. Thus, isolation of multiple cross-reactive pan-CoV S2 stem bnAb lineages, understanding their maturation pathways, and identifying common antibody framework motifs, are likely to be important for rational vaccine design approaches (*80*). Encouragingly, two recent studies have described a similar CoV S2 domain bnAb epitope being targeted by cross-reactive mAbs isolated from heterologous CoV spike protein immunizations in mice and mice transgenic for the human Ig locus (*52, 53*). These data suggest that such bnAbs could be induced by both immunization with designed vaccines as well as coronavirus infection in humans. Nonetheless, it would need to be ascertained how many sequential immunizations would be needed to broaden the breadth of these nAb responses.

Interestingly, despite relatively low neutralization potency, CC40.8 showed robust in vivo protective efficacy against SARS-CoV-2 challenge. The findings illustrate that extra-neutralizing effector functions of S2 stem bnAbs may be important and need to be investigated to determine their role in viral suppression or clearance. Indeed, recent studies have revealed that cross-reactive antibodies to endemic CoVs could serve as a marker for survival after severe disease (*81, 82*) or protection against COVID-19 (*83*).

Although we show CC40.8 S2 stem-helix bnAb confers in vivo protection against SARS-CoV-2 infection, our study has limitations. First, our CC40.8 protection studies were focused on SARS-CoV-2 infection and testing the protective efficacy of CC40.8 against a broad range of betacoronaviruses will be important. Second, we showed CC40.8 protects against SARS-CoV-2 in two small animal models and investigating its protective efficacy in non-human primate models is also eventually desirable. Third, CC40.8 bnAb showed remarkable protection even at very low antibody levels despite its low neutralization potency, hence warranting future studies to investigate the role of other factors such as antibody effector function that may contribute to CC40.8-mediated protection against coronaviruses.

Overall, we describe a cross-neutralizing human bnAb epitope on β-CoVs and provide molecular details that help explain its broad reactivity. The identification of this conserved epitope in the coronavirus spike protein should facilitate bnAb-epitope based vaccine development and antibody-based intervention strategies not only to SARS-CoV-2, but against existing human coronaviruses and other coronaviruses that could emerge with pandemic potential.

### Materials and Methods Study Design

The objective of the study was to evaluate a previously discovered SARS-CoV-2 spike protein stem-helix antibody, CC40.8, for binding to and neutralization of diverse sarbecoviruses and SARS-CoV-2 Variants of Concern, structurally define its epitope site and test its protective efficacy. For in vitro binding and neutralization studies, CC40.8 and control mAbs were tested in duplicate and experiments were repeated independently for rigor and reproducibility. We did not use any statistical methods to predetermine sample sizes for the animal studies. All hACE2-trangenic mouse or hamster experiments used 5 or 6 animals per group. A positive and/or a negative control mAb-treated animal group was included in the in vivo SARS-CoV-2 challenge experiments. Male and female, age matched (8-week old) animals were randomly assigned in CC40.8 bnAb-treated or control mAb-treated animal groups for the SARS-CoV-2 challenge studies. All immunological and virological measurements were performed blinded. Animals were euthanized at day five post infection to measure weight loss and lung viral load. The serum antibody titers of the passively transferred antibody were determined daily for the SARS-CoV-2 challenge experiment in the hACE2 mice. No datapoints were excluded as outliers in any experiment.

### Human cohort information

Plasma from convalescent COVID-19 donors were kindly provided through the “Collection of Biospecimens from Persons Under Investigation for 2019-Novel Coronavirus Infection to Understand Viral Shedding and Immune Response Study” UCSD IRB# 200236. Samples were collected based on COVID-19 diagnosis regardless of gender, race, ethnicity, disease severity, or other medical conditions. The gender for individuals was evenly distributed across the human cohort. All human donors were assessed for medical decision-making capacity using a standardized, approved assessment, and voluntarily gave informed consent prior to being enrolled in the study. The summary of the demographic information of the COVID-19 donors is listed in table S2.

### Pseudovirus production and generation of mutant spike proteins

Under biosafety level 2 and 3 conditions, MLV-gag/pol (Addgene #14887) and pCMV-Fluc (Addgene #170575) were co-transfected into HEK293T cells along with plasmids encoding full-length or variously truncated spike proteins from SARS-CoV-1, WIV1, SHC014, PANG17, MERS-CoV and SARS-CoV-2 (SARS-CoV-2 variants of concern (B.1.1.7 (alpha), B.1.351 (beta), P.1 (gamma) and B.1.617.2 (delta)) using Lipofectamine 2000 (Thermo Fisher Scientific cat.# 11668019) to produce single-round of infection competent pseudo-viruses. The media was changed by fresh Dulbecco’s Modified Eagle Medium (DMEM) with 10% heat-inactivated FBS, 4mM L-Glutamine and 1% P/S 16 hours post transfection. The supernatant containing MLV-pseudotyped viral particles was collected 48 hours post transfection, aliquoted and frozen at -80 °C for the neutralization assay. Amino-acid point mutations in SARS-CoV-2 spike protein-encoding plasmids were made by using site-directed mutagenesis kit (New England Biolabs cat.# E0554S) according to the manufacturer’s instructions. All the mutations were verified by DNA sequence analysis (Eton Bioscience).

### Neutralization assay

Pseudotyped viral neutralization assay was performed as previously described with minor modifications (Modified from TZM-bl assay protocol (*84*)). In sterile 96-well half-area plates (Corning cat.# 3688), 25 μl of virus was immediately mixed with 25 μl of three-fold serially diluted monoclonal antibodies (mAb) (starting concentration of 300 µg/ml) or serially diluted plasma from COVID-19 donors and incubated for one hour at 37°C to allow for antibody neutralization of the pseudotyped virus. Synthesized peptides were optionally added in the mixture for testing inhibition of neutralization. 10,000 HeLa-hACE2 cells (as previously generated (*33*)) per well (in 50 µl of media containing 20 μg/ml Dextran) were directly added to the antibody virus mixture. Plates were incubated at 37°C for 42 to 48 hours. Following the infection, HeLa-hACE2 cells were lysed using 1x luciferase lysis buffer (25mM Gly-Gly pH 7.8, 15mM MgSO4, 4mM EGTA, 1% Triton X-100). Luciferase intensity was then read on a Luminometer with luciferase substrate according to the manufacturer’s instructions (Promega cat.# E2620). Percentage of neutralization was calculated using the following equation: 100 X (1 – (mean fluorescent intensity (MFI) of sample – average MFI of background) / average of MFI of probe alone – average MFI of background)). Fifty percent maximal inhibitory concentrations (IC_50_), the concentrations required to inhibit infection by 50% compared to the controls, were calculated using the dose-response-inhibition model with 5-parameter Hill slope equation in GraphPad Prism 7 (GraphPad Software)

### Flow cytometry based Cellular-ELISA (CELISA) binding

Binding of monoclonal antibody to various human coronavirus (HCoV) spike proteins expressed on the surface of HEK293T cells was determined by flow cytometry, as described previously (*85*). Briefly, HEK293T cells were transfected with different plasmids encoding full-length HCoV spike proteins and were incubated for 36 to 48 hours at 37°C. Post incubation cells were trypsinized to prepare a single cell suspension and were distributed into 96-well plates. Monoclonal antibodies were prepared as 5-fold serial titrations in FACS buffer (1x phosphate-buffered saline (PBS), 2% fetal bovine serum (FBS), 1 mM EDTA), starting at 10 µg/ml, 6 dilutions. 50 μl/well of the diluted samples were added into the cells and incubated on ice for 1 hour. The plates were washed twice in FACS buffer and stained with 50 μl/well of 1:200 dilution of R-phycoerythrin (PE)-conjugated mouse anti-human IgG Fc antibody (SouthernBiotech cat.# 9040-09) and 1:1000 dilution of Zombie-NIR viability dye (BioLegend cat.# 423105) on ice in dark for 45 minutes. After another two washes, stained cells were analyzed using flow cytometry (BD Lyrics cytometers), and the binding data were generated by calculating the percent (%) PE-positive cells for antigen binding using FlowJo 10 software.

### Expression and purification of HCoV spike proteins and SARS-CoV-2 spike protein mutants

To express the soluble spike ectodomain proteins, the HCoV spike protein encoding plasmids were transfected into FreeStyle293-F cells (Thermo Fisher Scientific cat.# R79007). For general production, 350 µg of plasmids were transfected into 1L FreeStyle293-F cells at the density of 1 million cells per mL. 350 µg plasmids in 16 ml Opti-MEM™ (Thermo Fisher Scientific cat.# 31985070) were filtered and mixed with 1.8 mL 40K PEI (1mg/mL) in 16 ml Opti-MEM™. After gently mixing the two components, the combined solution rested at room temperature for 30 minutes and was poured into 1L FreeStyle293-F cell culture. The cell cultures were centrifuged at 2500xg for 15 minutes on day 4 after transfection, and the supernatants were filtered through the 0.22 μm membrane. The His-tagged proteins were purified with the HisPur Ni-NTA Resin (Thermo Fisher Scientific cat.# 88221). Each column was washed with at least 3 bed volumes of wash buffer (25 mM Imidazole, pH 7.4), followed by elution with 25 ml of the elution buffer (250 mM Imidazole, pH 7.4) at slow gravity speed (about 4 seconds per drop). The eluates were buffer exchanged into PBS by using Amicon tubes, and the proteins were concentrated afterwards. The proteins were further purified by size-exclusion chromatography using a Superdex 200 Increase 10/300 GL column (GE Healthcare cat.# GE28-9909-44). The selected fractions were pooled and concentrated again for further use.

### BioLayer Interferometry (BLI) binding

The determination of monoclonal antibody binding with spike proteins or selected peptides was conducted in an Octet K2 system (ForteBio). The anti-human IgG Fc capture (AHC) biosensors (ForteBio cat.# 18-5063) were used to capture IgG first for 60 seconds. After providing baseline in Octet buffer for another 60 seconds, the sensors were transferred into HCoV spike proteins at various concentrations for 120 seconds for association, and into Octet buffer for disassociation for 240 seconds. Alternatively, the hydrated streptavidin biosensors (ForteBio cat.# 18-5020) first captured the N-terminal biotinylated peptides diluted in Octet buffer (PBS plus 0.1% Tween-20) for 60 seconds, then transferred into Octet buffer for 60 seconds to remove unbound peptide and provide the baseline. Then the sensors were immersed in diluted monoclonal antibody IgG for 120 seconds to provide association signal, followed by transferring into Octet buffer to test for disassociation signal for 240 seconds. The data generated was analyzed using the ForteBio Data Analysis software for correction, and the kinetic curves were fit to 1:1 binding mode. Note that the IgG: spike protomer binding can be a mixed population of 2:1 and 1:1, such that the term ‘apparent affinity’ dissociation constants (KD^App^) are shown to reflect the binding affinity between IgGs and spike trimers tested.

### HEp2 epithelial cell polyreactive assay

Reactivity to human epithelial type 2 (HEp2) cells was determined by indirect immunofluorescence on HEp2 slides (Hemagen, cat.# 902360) according to manufacturer’s instructions. Briefly, monoclonal antibody was diluted at 50 μg/mL in PBS and then incubated onto immobilized HEp2 slides for 30 minutes at room temperature. After washing 3 times with PBS buffer, one drop of fluorescein isothiocyanate (FITC)-conjugated goat anti-human IgG was added onto each well and incubated in the dark for 30 minutes at room temperature. After washing, the coverslip was added to HEp2 slide with glycerol and the slide was photographed on a Nikon fluorescence microscope to detect FITC signal.

### Polyspecificity reagent (PSR) ELISA

Solubilized CHO cell membrane protein (SMP) was coated onto 96-well half-area high-binding ELISA plates (Corning, cat.# 3690) overnight at 4°C. After washing with PBS plus 0.05% Tween-20 (PBST), plates were blocked with 3% bovine serum albumin (BSA) for 2 hours at 37°C. Antibody samples were diluted at 10 μg/mL in 1% BSA with 3-fold serial dilution and then added in plates to incubate for 1 hour at 37°C. After 3 thorough washes with PBST, alkaline phosphatase-conjugated goat anti-human IgG Fc secondary antibody (Jackson ImmunoResearch, cat.# 109-055-008) was added to the plate and incubated for 1 hour at 37°C. After a final wash, phosphatase substrate (Sigma-Aldrich, cat.# S0942-200TAB) was added into each well. Absorption was measured at 405 nm.

### Peptide scanning by ELISA binding

N-terminal biotinylated overlapping peptides corresponding to the complete sequence of HCoV-HKU1 S2 subunit (residue number range: 761-1355 (GenBank: AAT98580.1) were synthesized at A&A Labs (Synthetic Biomolecules). Each peptide was 15 residue long with a 10 amino acid overlap. For ELISA binding, 96-well half-area plates (Corning cat. # 3690) were coated overnight at 4°C with 2 µg/ml of streptavidin in PBS. Plates were washed 3 times with PBST and blocked with 3% (wt/vol) BSA in PBS for 1 hour. After removal of the blocking buffer, the plates were incubated with peptides in 1% BSA plus PBST for 1.5 hours at room temperature. After a washing step, monoclonal antibody or serum samples diluted in 1% BSA/PBST were added into each well and incubated for 1.5 hours. DEN3 human antibody was used as a negative control. After the washes, a secondary antibody conjugated with alkaline phosphatase-conjugated goat anti-human IgG Fc secondary antibody (Jackson ImmunoResearch, cat.# 109-055-008) diluted 1:1000 in 1% BSA/PBST, was added to each well and incubated for 1 hour. The plates were then washed and developed using alkaline phosphatase substrate pNPP tablets (Sigma-Aldrich, cat.# S0942-200TAB) dissolved in stain buffer. The absorbance was recorded at an optical density of 405 nm (OD405) using a VersaMax microplate reader (Molecular Devices), where data were collected using SoftMax software version 5.4.

### Cell-cell fusion inhibition assay

HeLa stable cell lines were generated through transduction of lentivirus carrying genes encoding either human ACE2 (hACE2) and enhanced green fluorescent protein (EGFP) or nuclear localization signal (NLS)-mCherry and SARS-CoV-2 spike protein. The pBOB construct carrying these genes was co-transfected into HEK293T cells along with lentiviral packaging plasmids pMDL, pREV, and pVSV-G (Addgene #12251, #12253, #8454) by Lipofectamine 2000 (Thermo Fisher Scientific, cat.# 11668019) according to the manufacturer’s instructions. Supernatants were collected 48 hours after transfection, then were transduced to pre-seeded HeLa cells. 12 hours after transduction, stable cell lines were collected, scaled up and stored for cell-cell fusion assay. 10,000 NLS-mCherry^+^ HeLa cells expressing SARS-COV-2 spike protein were seeded into 96-well half-well plates on the day before the assay. The culture medium was removed by aspiration before the assay. 50 µl of 50 µg/ml CC40.8 and DEN3 mAbs diluted in DMEM with 10% heat-inactivated FBS, 4mM L-Glutamine and 1% P/S were then added to the pre-seeded cells and incubated for 1 hour in an incubator. 50 µl of 10,000 EGFP^+^hACE2^+^ HeLa cells were added to the plates and incubated for 2 hours before taking images under the microscope.

### Sequence alignments of coronavirus spike stem regions

The spike sequences of SARS-CoV-2, SARS-CoV-1, RaTG13, SHC014, Rs4081, Pang17, RmYN02, Rf1, WIV1, Yun11, BM48-31, BtKY72, HCoV-HKU1, HCoV-OC43, MERS-CoV, MHV, HCoV-229E and HCoV-NL63 were downloaded from the GenBank and aligned against the SARS-CoV-2 reference sequence using BioEdit (http://www.mbio.ncsu.edu/bioedit/bioedit.html).

### Expression and purification of CC40.8 Fab

To generate Fab, CC40.8 IgG was digested by Papain (Sigma-Aldrich cat.# P3125) for 4 hours at 37 °C, then was incubated with Protein-A beads at 4 °C for 2 hours to remove the Fc fragments. CC40.8 Fab was concentrated afterwards and further purified by size-exclusion chromatography using a Superdex 200 Increase 10/300 GL column (GE Healthcare cat.# GE28-9909-44). The selected fractions were pooled and concentrated again for further use.

### Crystallization and structural determination

A mixture of 9 mg/ml of CC40.8 Fab and 10× (molar ratio) SARS-CoV-2 stem peptide was screened for crystallization using the 384 conditions of the JCSG Core Suite (Qiagen) on our robotic CrystalMation system (Rigaku) at Scripps Research. Crystallization trials were set-up by the vapor diffusion method in sitting drops containing 0.1 μl of protein and 0.1 μl of reservoir solution. Optimized crystals were then grown in drops containing 0.1 M sodium acetate buffer at pH 4.26, 0.2 M ammonium sulfate, and 28% (w/v) polyethylene glycol monomethyl ether 2000 at 20°C. Crystals appeared on day 7, were harvested on day 15 by soaking in reservoir solution supplemented with 20% (v/v) glycerol, and then flash cooled and stored in liquid nitrogen until data collection. Diffraction data were collected at cryogenic temperature (100 K) at Stanford Synchrotron Radiation Lightsource (SSRL) on the Scripps/Stanford beamline 12-1, with a beam wavelength of 0.97946 Å, and processed with HKL2000 (*86*). Structures were solved by molecular replacement using PHASER (*87*). A model of CC40.8 was generated by Repertoire Builder (https://sysimm.ifrec.osaka-u.ac.jp/rep_builder/) (*88*). Iterative model building and refinement were carried out in COOT (*89*) and PHENIX (*90*), respectively. Epitope and paratope residues, as well as their interactions, were identified by accessing PISA at the European Bioinformatics Institute (http://www.ebi.ac.uk/pdbe/prot_int/pistart.html) (*91*).

### Animal Study

8-week old transgenic hACE2 mice were given an i.p. antibody injections 12 hours pre-infection. Mice were infected through intranasal installation of 2X10^4^ total plaque-forming units (PFU) per animal of SARS-CoV-2 (USA-WA1/2020) in 25 µL of DMEM. Mice were bled on days 1, 2, 3, and 5 for serum antibody detection and weighed for the duration of the study. At day 5 post-infection, animals were euthanized, and lungs were harvested for quantitative polymerase chain reaction (qPCR) viral titer analysis and plaque live virus analysis. Similar experimental procedures were conducted for the protection study in 8-week old Syrian hamsters except that a higher SARS-CoV-2 (USA-WA1/2020) challenge dose (10^6^ total PFU per animal) was used. The research protocol was approved and performed in accordance with Scripps Research IACUC Protocol #20-0003

### Antibody detection in hACE2 serum samples by ELISA

Serum samples were obtained at day 1, 2, 3, and 5 to quantify mAb titers. Unconjugated F(ab’)₂ fragment of goat anti-human F(ab’)₂ fragment (Jackson ImmuoResearch cat.# 109-006-097) was coated to the ELISA plates overnight, then washed by PBS plus 1% Tween-20 three times. After being blocked by 3% BSA for 2 hours at 37°C, mouse serum dilution series and CC40.8 mAb dilution series for a standard curve were applied to the plates and reacted for 1 hour at 37°C. After three thorough washes with PBS plus 1% Tween-20, alkaline phosphatase-conjugated goat anti-human IgG Fc secondary antibody (Jackson ImmunoResearch, cat.# 109-055-008) was added to the plates before washing 3 times with PBS plus 1% Tween-20 and AP substrate applied for detection. The plates were read at 405nm and data were analyzed by CurveExpert.

### SARS-CoV-2 RNA Quantification

Viral RNA was isolated from lung tissue and subsequently amplified and quantified in a reverse transcription (RT)-qPCR reaction. Lung tissue was extracted at day 5 post infection and placed in 1 mL of trIzol reagent (Invitrogen). The samples were then homogenized using a Bead Ruptor 12 (Omni International). Tissue homogenates were then spun down and the supernatant was added to an RNA purification column (Qiagen). Purified RNA was eluted in 60 μL of DNase-, RNase-, endotoxin-free molecular biology grade water (Millipore). RNA was then subjected to reverse transcription and quantitative PCR using the CDC’s N1 (nucleocapsid) primer sets (Forward 5’-GAC CCC AAA ATC AGC GAA AT-3’; Reverse 5’-TCT GGT TAC TGC CAG TTG AATCTG-3’) and a fluorescently labeled (FAM) probe (5’-FAM-ACC CCG CAT TAC GTT TGGTGG ACC-BHQ1-3’) (Integrated DNATechnologies) on a BioRad CFX96 Real-Time instrument. For quantification, a standard curve was generated by diluting 2.5X10^6^ PFU RNA equivalents of SARS-CoV-2. Every run utilized eleven 5-fold serial dilutions of the standard. SARS-CoV-2-negative mouse lung RNA and no templates were both included as negative controls for the extraction step as well as the qPCR reaction.

### Viral load measurements

SARS-CoV-2 titers were measured by homogenizing organs in DMEM plus 2% fetal calf serum using 100 µm cell strainers (Myriad cat.# 2825-8367). Homogenized organs were titrated 1:10 over six steps and layered over Vero-E6 cells. After 1 hour of incubation at 37°C, a 1% methylcellulose in DMEM overlay was added, and the cells were incubated for 3 days at 37°C. Cells were fixed with 4% paraformaldehyde and plaques were counted by crystal violet staining.

### Statistical Analysis

Statistical analysis was performed using Graph Pad Prism 8 for Mac (Graph Pad Software). Groups of data were compared using the Kruskal-Wallis non-parametric test. Dunnett’s multiple comparisons test were also performed between experimental groups. Data were considered statistically significant at p < 0.05.

## Acknowledgements

We thank all of the human cohort participants for donating samples.

## Author contributions

P.Z., M.Y., G.S., I.A.W., D.R.B., and R.A. conceived and designed the study. N.B., J.R., M.P., E.G., S.A.R., D.M.S., and T.F.R. recruited donors and collected and processed plasma samples. P.Z., G.S., and F.A., performed BLI, ELISA and cell binding and virus neutralization assays. D.H., and L.P., conducted cell-cell fusion experiment. W.H., S.C., and P.Y., generated recombinant protein antigens. M.Y. and X.Z. determined the crystal structure of the antibody-antigen complex. N.B., N.S., J.R.T., and T.F.R. carried out animal studies and viral load measurements. P.Z., M.Y., G.S., N.B., N.S., D.H., W.H., D.N., J.R.T., T.F.R., I.A.W., D.R.B., and R.A. designed the experiments and analyzed the data. R.A., P.Z., G.S., M.Y., I.A.W. and D.R.B. wrote the paper, and all authors reviewed and edited the paper.

## Data Availability

All data associated with this study are in the paper or supplementary materials.

## SUPPLEMENTARY MATERIALS

**Fig. S1.**
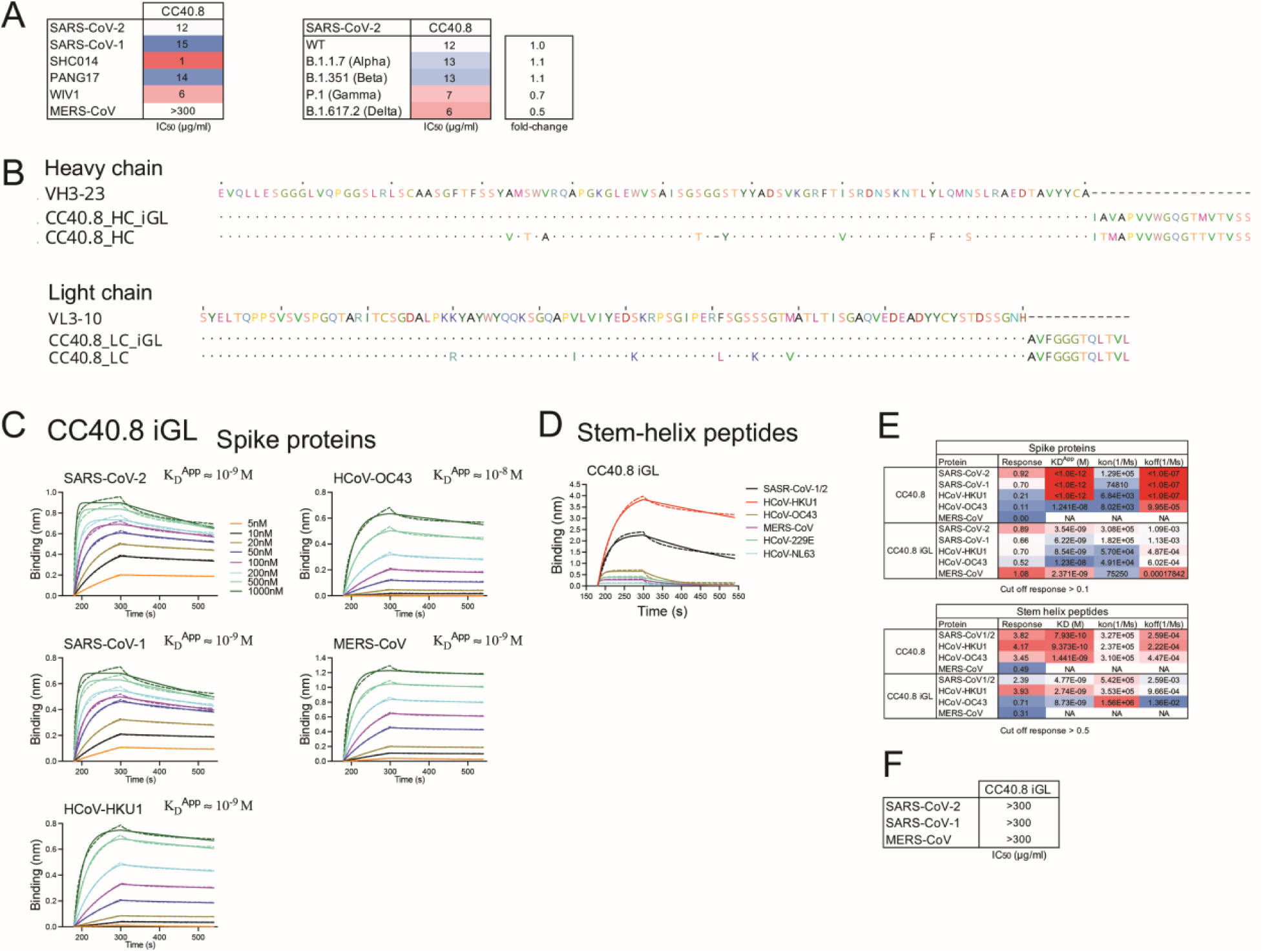
CC40.8 mature and CC40.8 iGL antibodies bind to spike proteins and stem-helix peptides and mature antibody neutralizes pseudotyped coronaviruses. **(A)** IC50 neutralization of CC40.8 broadly neutralizing antibody (bnAb) is shown for sarbecoviruses (SARS-CoV-2, SARS-CoV-1, SHC014, Pang17 and WIV1), MERS-CoV and SARS-CoV-2 variants of concern (alpha (B.1.1.7), beta (B.1.351), gamma (P.1) and delta (B.1.617.2)). **(B)** Sequence alignment of CC40.8 heavy and light chains with their corresponding germline V-gene sequences (VH3-23 and VL3-10) is shown with the design of CC40.8 antibody inferred germline (iGL) gene sequences. Dots represent identical residues and dashes represent gaps introduced to preserve the alignment. **(C)** BioLayer Interferometry (BLI) binding is shown for CC40.8 iGL antibody with human β-HCoV soluble spike proteins. Apparent binding constants (KD^App^) for each antibody-antigen interaction are indicated. The raw experimental curves are shown as dash lines, while the solid lines are the fits. **(D)** BLI binding is shown for CC40.8 iGL Ab to 25-mer stem peptides derived from all HCoV spike proteins. The kinetic curves are fit with a 1:1 binding mode. **(E)** Binding kinetics (KD^App^ (spike proteins), KD (stem-helix peptides) *kon* and *koff* constants) of CC40.8 and CC40.8 iGL antibodies with human β-HCoV soluble spike proteins and the 25-mer β-HCoV stem peptides are shown. **(F)** IC50 neutralization of CC40.8 iGL is shown for sarbecoviruses (SARS-CoV-2 and SARS-CoV-1) and MERS-CoV.

**Fig. S2.**
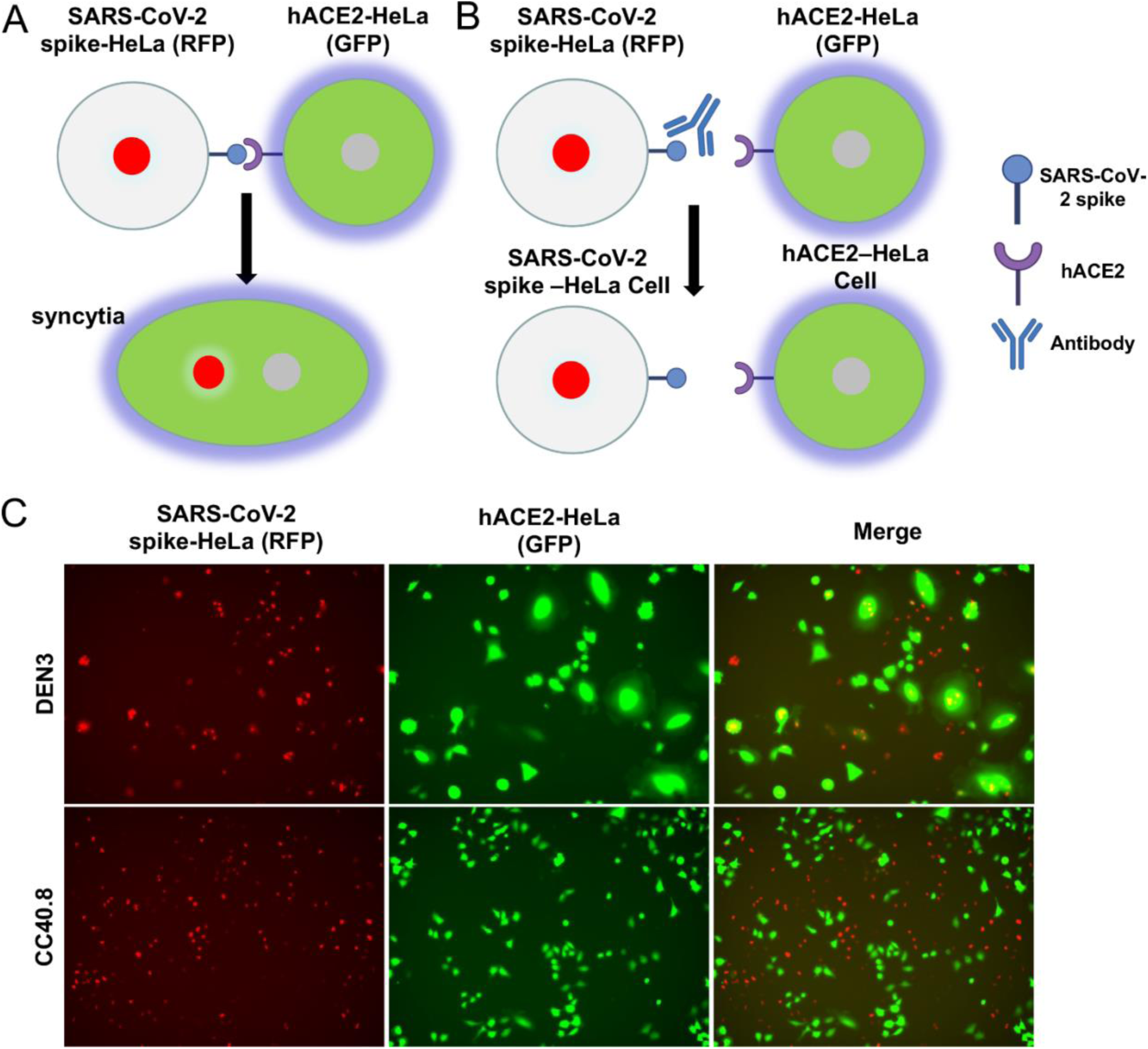
CC40.8 antibody inhibits SARS-CoV-2 spike protein- and hACE2-mediated cell-cell fusion. **(A and B)** A schematic diagram of cell-cell fusion assay is shown. SARS-CoV-2 spike-HeLa cells express nucleus-restricted RFP (Red) and hACE2-HeLa cells express cytosolic GFP (Green). The interaction of SARS-CoV-2 spike protein and hACE2 can lead to cell fusion to form syncytia. In the same syncytium, both GFP in the cytoplasm and RFP in the nucleus can be seen (A). If antibody can block cell-cell fusion, no syncytia can be seen. Only GFP-expressing hACE2-HeLa cells and RFP-expressing SARS-CoV-2 spike-HeLa cells can be seen (B). **(C)** SARS-CoV-2 spike-HeLa cells (red) were pre-incubated with negative control antibody (DEN3) or CC40.8 S2 stem bnAb for 1 hour, and then mixed with hACE2-HeLa cells (green). Green syncytia were observed with DEN3, indicating widespread cell-cell fusion mediated by SARS-CoV-2 spike and hACE2; fusion was inhibited by addition of CC40.8.

**Fig. S3.**
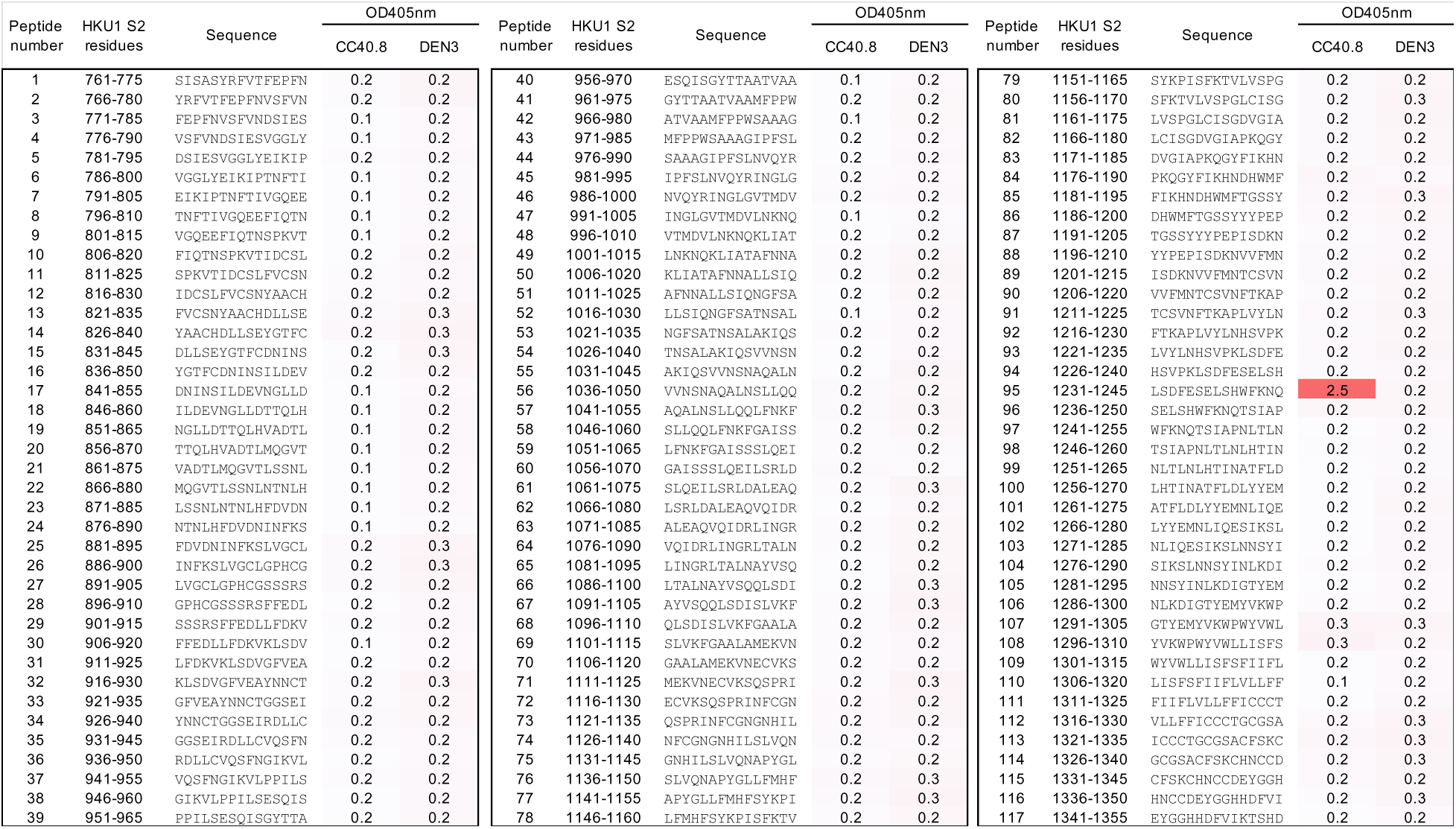
**Epitope mapping of CC40.8 antibody with HCoV-HKU1 S2 subunit-derived overlapping peptides.** ELISA binding results are shown for CC40.8 mAb with HCoV-HKU1 S2 subunit overlapping peptides (residue number range: 761-1355). Each HCoV-HKU1 S2 subunit peptide is 15-residues long with a 10-residue overlap. Peptide IDs, S2 subunit residue number ranges of 15-mer peptides, and antibody binding responses are shown. CC40.8 exhibited binding to the 95^th^ peptide (residue position range: 1231-1245) corresponding to the HCoV-HKU1 S2 stem-helix region. DEN3, an antibody to dengue virus, was used as a control.

**Fig. S4.**
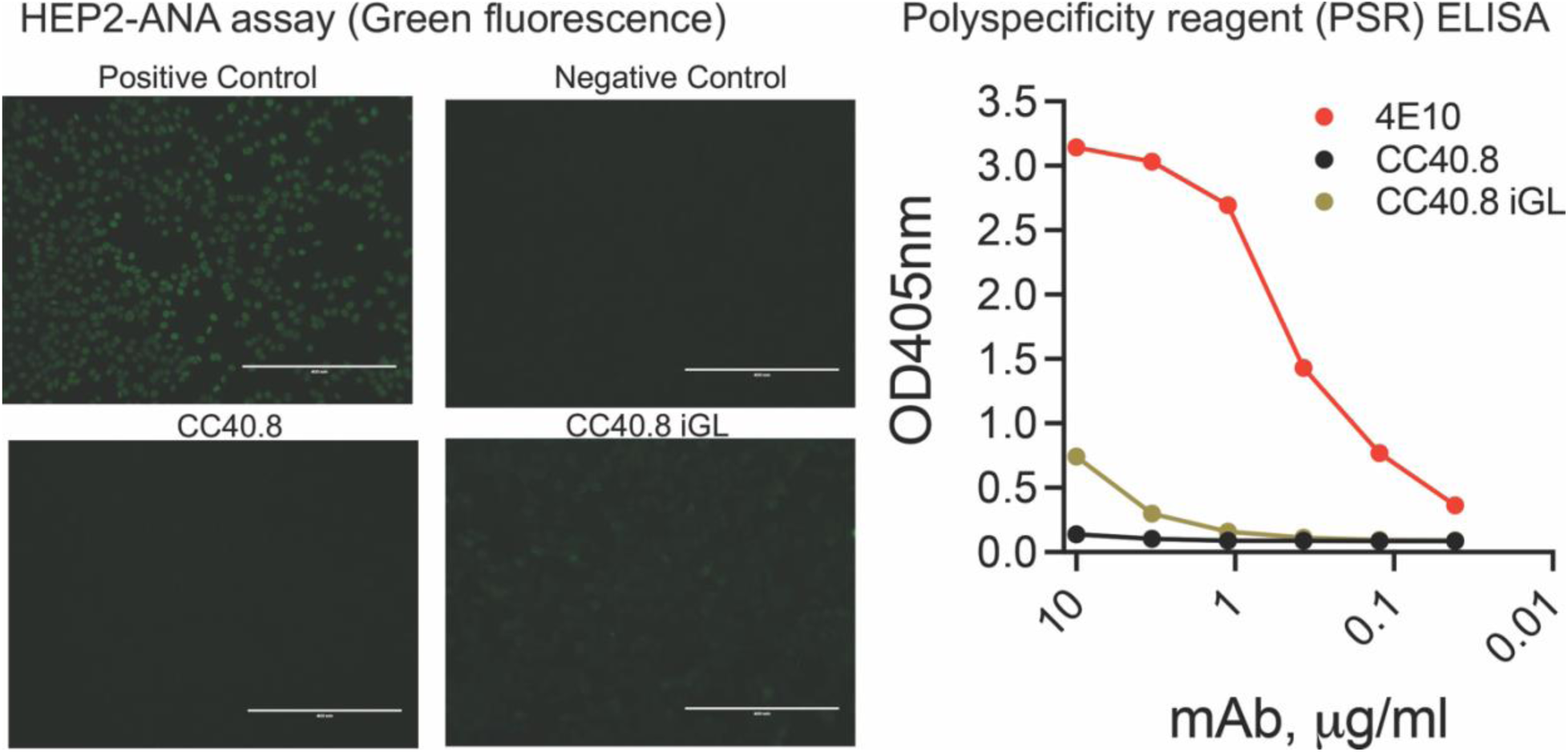
Polyreactivity analysis of CC40.8 and CC40.8 iGL antibodies. **(A)** Immunofluorescence showing binding of antibodies to immobilized HEp2 epithelial cells was detected by FITC-labelled secondary antibody. Positive and negative controls for the Hep2 kit assay are provided by the manufacturer. **(B)** Antibodies were tested by enzyme-linked immunosorbent assay (ELISA) for binding to the polyspecificity reagent (PSR) from CHO-cell solubilized membrane protein (SMP). 4E10, an HIV MPER-specific antibody known to display polyreactivity, was used as a positive control.

**Fig. S5.**
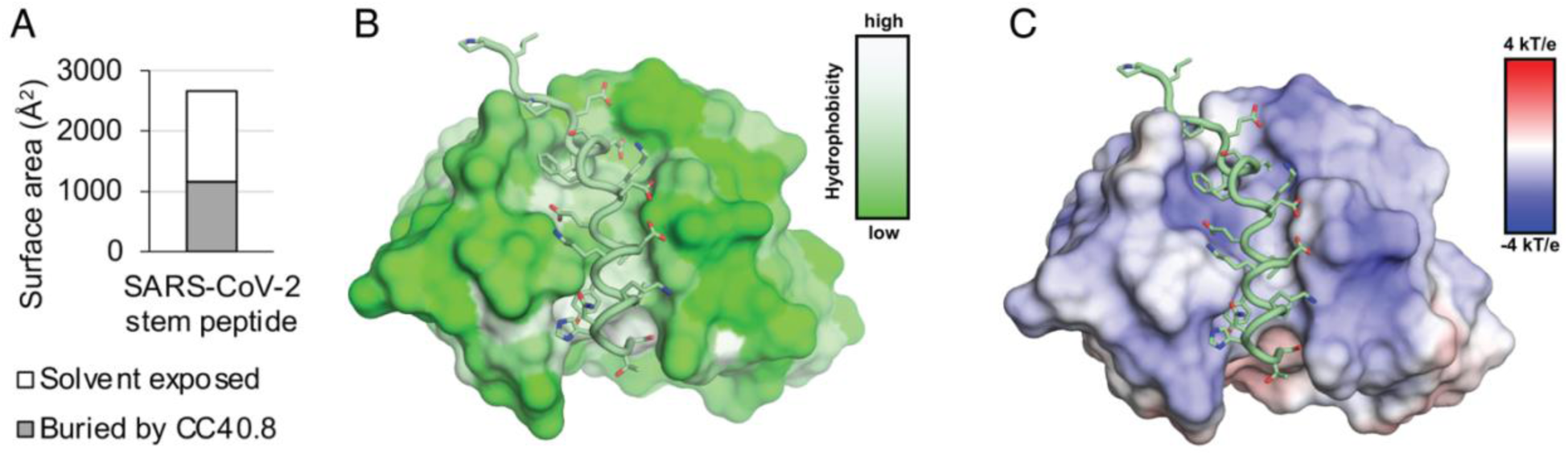
CC40.8 bnAb structure with SARS-CoV-2 S2 stem-helix peptide. **(A)** Surface area of the SARS-CoV-2 stem peptide is shown. Solvent exposed and buried areas were calculated with Proteins, Interfaces, Structures and Assemblies (PISA) (*91*). **(B)** The SARS-CoV-2 stem peptide inserts into a hydrophobic groove formed by the heavy and light chains of CC40.8. Surfaces of CC40.8 are color-coded by hydrophobicity [calculated by Color h (https://pymolwiki.org/index.php/Color_h)]. **(C)** Electrostatic surface potential of the CC40.8 paratope is shown. Electrostatic potential was calculated by APBS and PDB2PQR (*93, 94*).

**Fig. S6.**
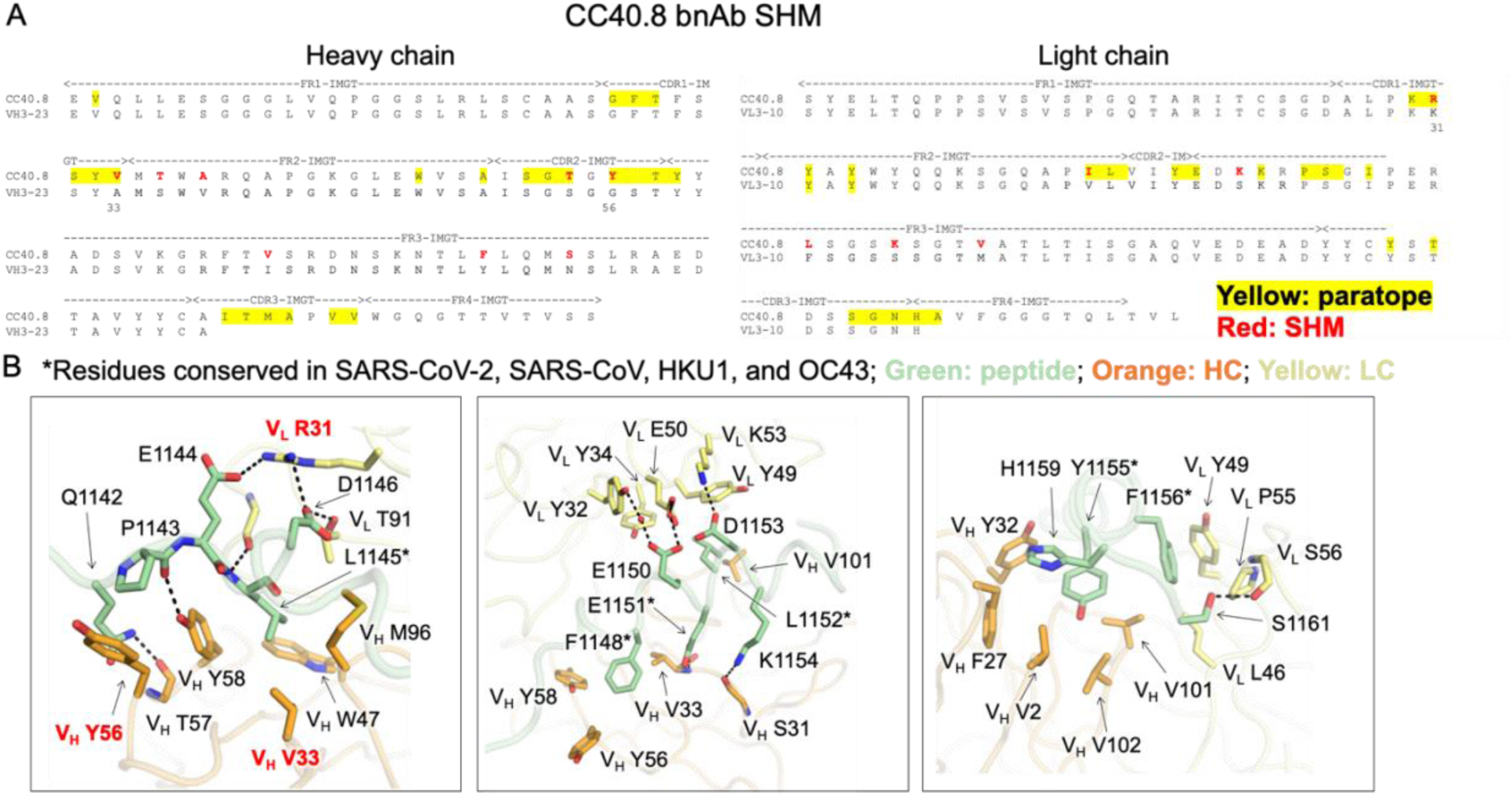
Contribution of CC40.8 bnAb heavy and light chain germline and somatic mutated V-gene residues in S2 stem epitope recognition. **(A)** Alignment of CC40.8 with germline VH3-23 and VL3-10 sequences is shown. Paratope residues [defined as buried surface area (BSA) > 0 Å^2^ as calculated by PISA (*91*)] are highlighted with yellow boxes. Somatically mutated residues as calculated by IgBLAST (*95*) are highlighted in red. B. Detailed interactions between CC40.8 Fab and the SARS-CoV-2 stem peptide are shown. Heavy and light chains of CC40.8 are shown in orange and yellow, while the SARS-CoV-2 stem peptide is in pale green. Hydrogen bonds and salt bridges are represented by black dashed lines. Somatically mutated residues are shown in red. Conserved residues among coronaviruses are indicated by asterisks (*).

**Fig. S7.**
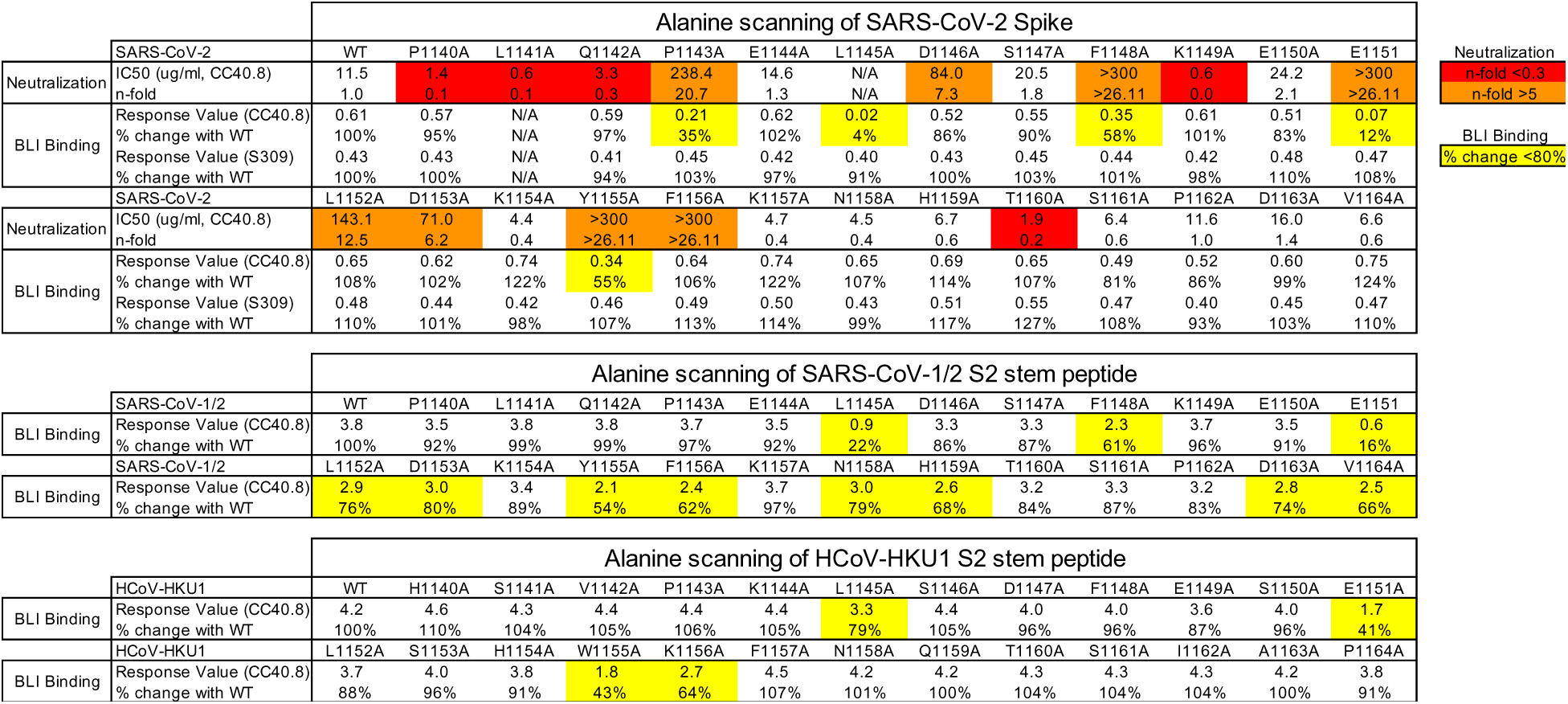
Epitope mapping of CC40.8 bnAb by alanine scanning mutagenesis of SARS-CoV-2 spike protein and SARS-CoV-2/HCoV-HKU1 S2 stem peptides using neutralization and BLI binding assays. The upper panel shows the IC50 neutralization of CC40.8 bnAb with wild-type (WT) SARS-CoV-2 and spike mutant pseudoviruses and the BLI binding responses with WT SARS-CoV-2 soluble spike protein and alanine mutants. SARS-CoV-2 receptor binding domain (RBD) antibody S309 was a control for the spike protein binding assays. The IC50 fold change (n-fold) was calculated by dividing the mutant value by the WT value. For IC50, n-fold <0.3 are indicated in red, and n-fold >5 in orange. The middle and lower panels show BLI binding responses of CC40.8 antibody to WT and alanine mutants of the SARS-CoV-1/2 and HCoV-HKU1 stem-helix peptides, respectively. Binding response values where the % change in binding (from WT peptide) is below 80% are indicated in yellow. Antibody S309 that recognizes a fairly conserved epitope of the RBD of both SARS-CoV-1 and SARS-CoV-2 was used as control. N/A, not available.

**Fig. S8.**
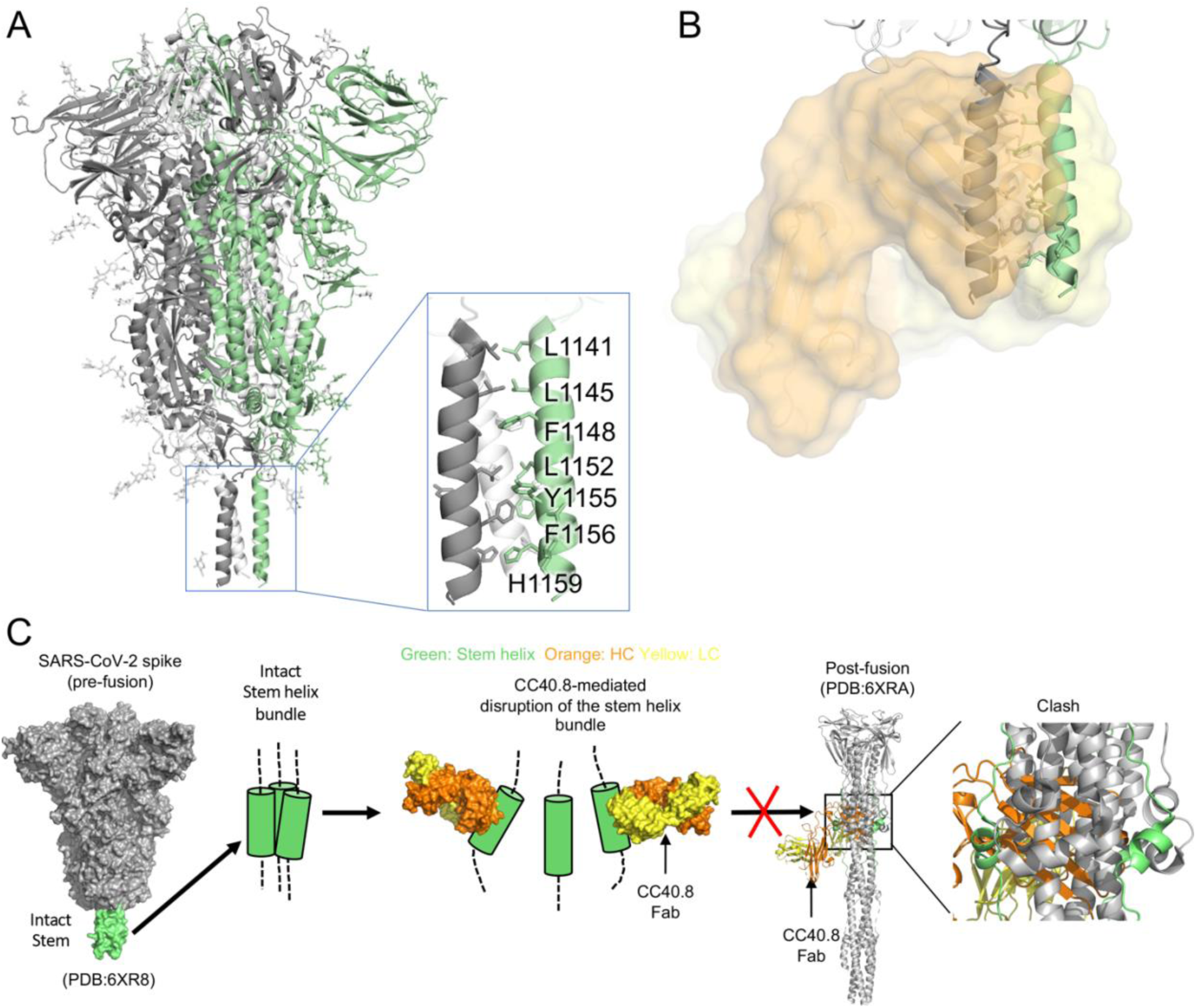
CC40.8 binds to a buried interface of the 3-helix bundle: predicted mechanism of neutralization. **(A)** A SARS-CoV-2 spike protein structure is shown in the pre-fusion state. The three protomers are shown in gray, pale green, and white, respectively, with N-linked glycans represented by sticks. The 3-helix bundle stem region is highlighted in a blue-outlined box. Representative epitope residues of CC40.8 are shown in sticks. The CC40.8 epitope is rich in hydrophobic residues. A cryo-EM structure of SARS-CoV-2 spike protein structure in the pre-fusion state that incorporates the coordinates of the 3-helix bundle stem region (PDB: 6XR8, (*96*)) is shown here. **(B)** The SARS-CoV-2 spike protein pre-fusion structure was superimposed on the structure of CC40.8 (orange/yellow) in complex with a SARS-CoV-2 S2 peptide. CC40.8 would clash with the other protomers of the spike protein in the pre-fusion state. **(C)** A putative neutralization mechanism of CC40.8 is presented. The S2 3-helix bundle region is shown in green, and heavy and light chains of CC40.8 are shown in orange and yellow, respectively. A model for the mechanism of neutralization is shown and inspired by the interaction of a mouse S2 stem antibody, B6, isolated from a spike protein vaccinated animal that targets a similar stem epitope (*52*) .

**Fig. S9.**
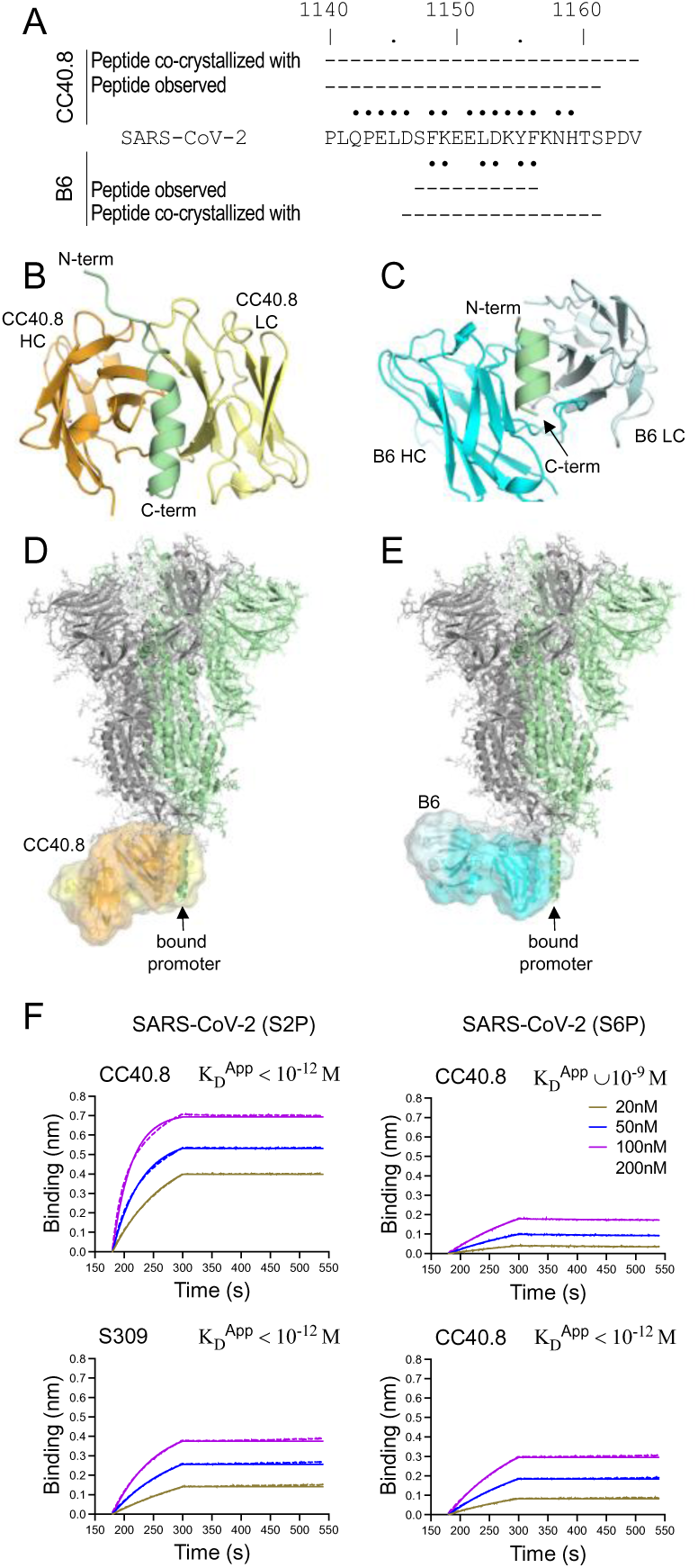
Comparison of bnAbs CC40.8 and B6 that target the S2 stem helix. **(A)** A comparison between S2 stem-helix peptides targeted by CC40.8 and B6 is shown. Peptides used for co-crystallization with CC40.8 or B6 are indicated by dashes, with the regions observed in the crystal structures of each study indicated. Residues involved in interactions with CC40.8 and B6 are indicated by black dots (cutoff distance = 4 Å). **(B to E)** Structures of CC40.8 and B6 were compared. The heavy and light chains of CC40.8 are colored in orange and yellow, respectively, and those for B6 are in cyan and light cyan. The S2 stem-helix peptides are shown in green. In panels (D) and (E), a SARS-CoV-2 spike protein pre-fusion structure (PDB 6XR8) is superimposed on structures of CC40.8 and B6 in complex with a SARS-CoV-2 S2 peptide in the green protomer (indicated by arrows). Both CC40.8 and B6 would clash with the other protomers of the spike protein in pre-fusion state. **(F)** BLI binding kinetics of CC40.8 to S2- and S6-stabilized SARS-CoV-2 spike trimers are shown. An RBD-targeting neutralizing Ab S309 (*97*) was used as a control. Apparent binding constants (KD^App^) of antibodies with spike proteins are shown. The raw experimental curves are shown as dash lines, while the solid lines are the fits.

**Figure S10.**
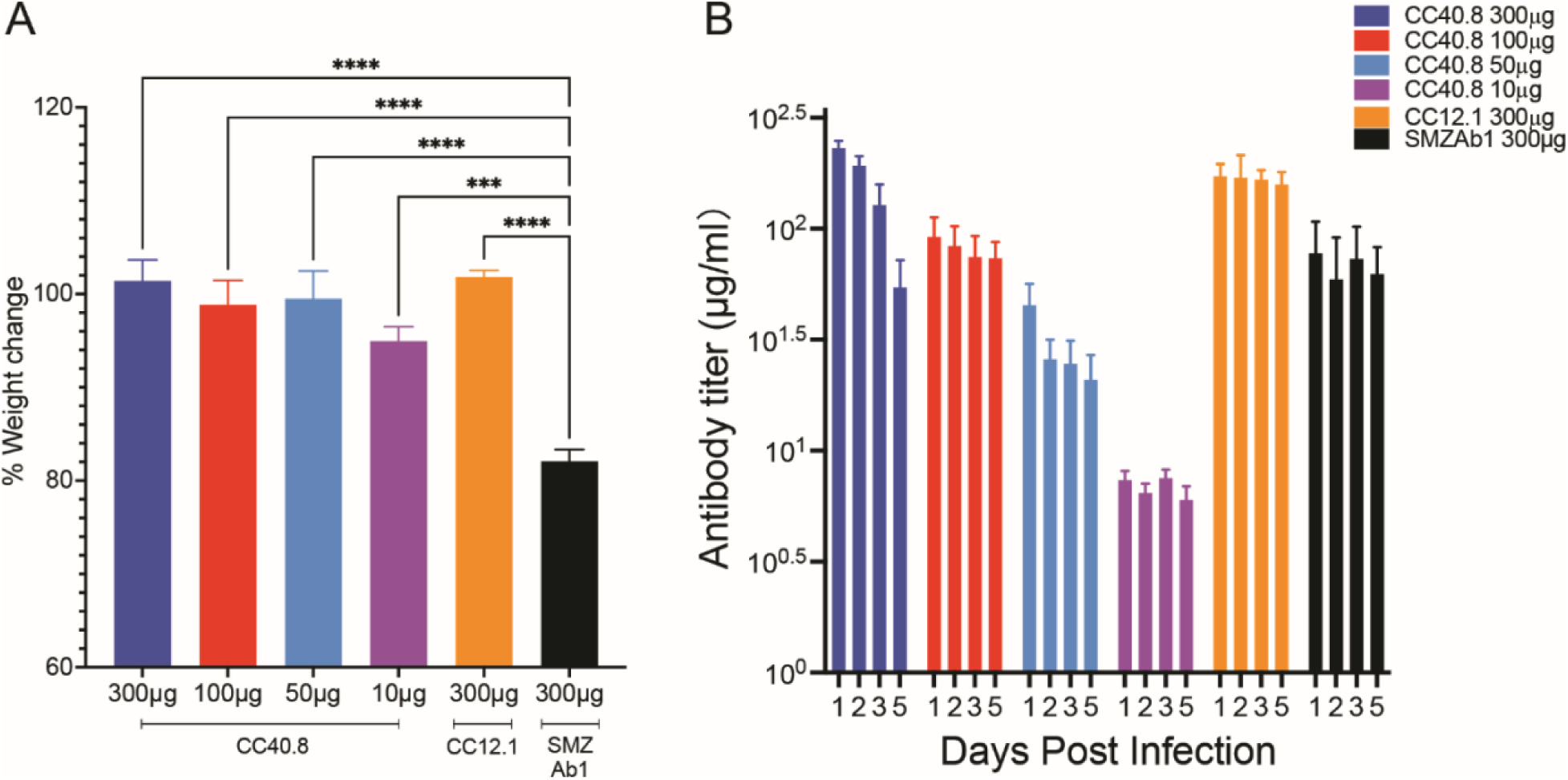
Weight loss, viral titers, and serum antibody titers were measured in hACE2 mice passively administered CC40.8. **(A)** Percent day 5 weight change was calculated from day 0 for all animals. Data are presented as mean ± SEM. Significance was calculated with Dunnett’s multiple comparisons test between each experimental group and the ZIKV Ab (SMZAb1) control group (***P<0.001, ****P<0.0001). **(B)** Serum human IgG concentrations of CC40.8, CC12.1 and SMZAb1 were assessed by ELISA at day 1, 2, 3, and 5 post infection. Data are presented as mean ± SEM.

**Fig. S11.**
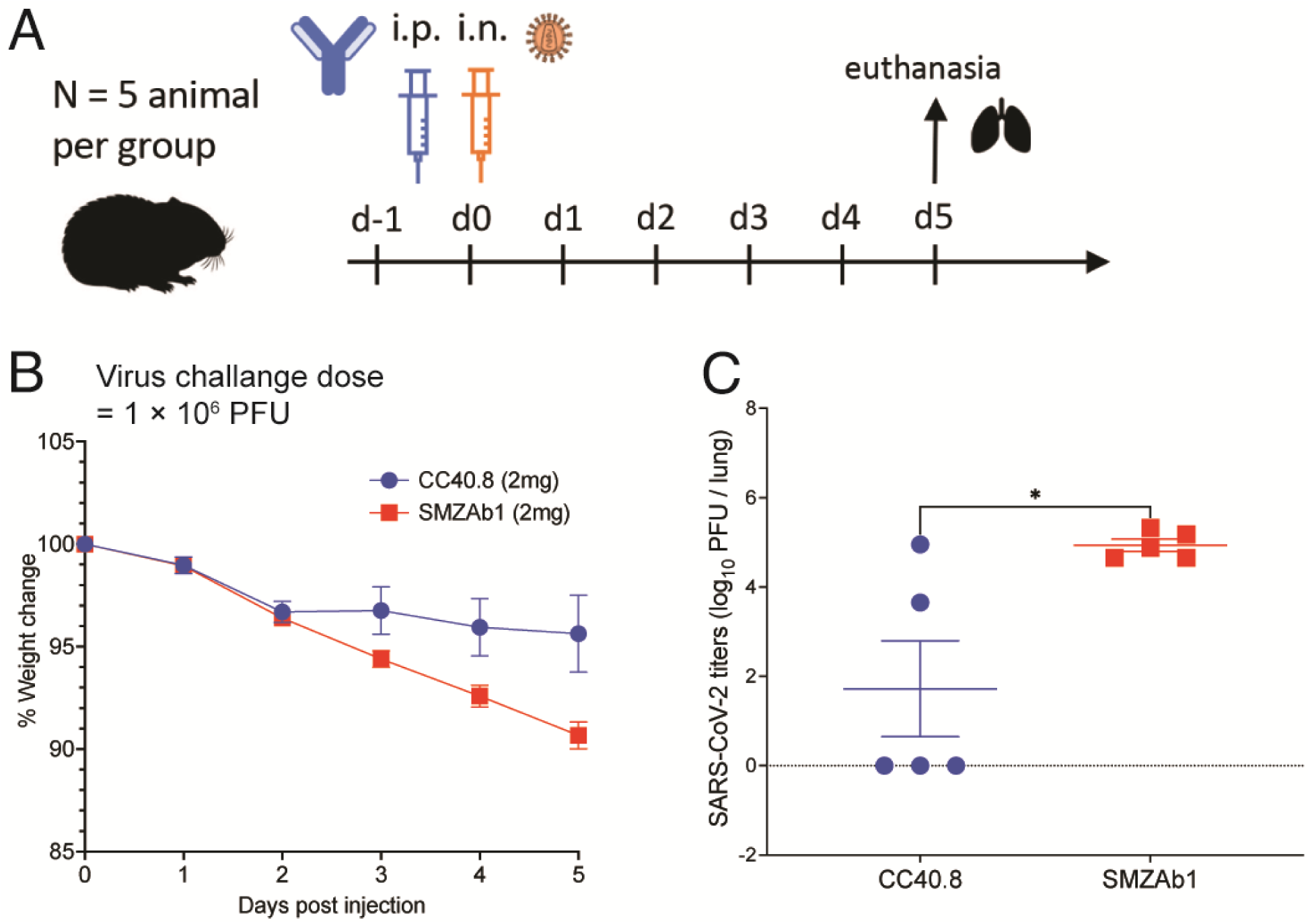
CC40.8 reduces weight loss and lung viral load and viral replication following SARS-CoV-2 challenge in Syrian hamsters. **(A)** CC40.8 was administered intraperitonially (i.p.) at a 2 mg per animal dose into Syrian hamsters (average: 16.5 mg/kg). Control animals received 2 mg of control SMZAb1. Each group of five animals was challenged intranasally (i.n.) 12 hours after antibody infusion with 1 × 10^6^ PFU of SARS-CoV-2. Animal weight was monitored daily as an indicator of disease progression and lung tissue was collected on day 5 for viral burden assessment. **(B)** Percent weight change is shown for CC40.8 or control antibody-treated animals after SARS-CoV-2 challenge. Percent weight change was calculated from day 0 for all animals. Data are presented as mean ± SEM. **(C)** SARS-CoV-2 titers (PFU) were determined by plaque assay from lung tissue at day 5 after infection. Three out of 5 CC40.8-treated animals had substantially reduced viral titers compared to the SMZAb1 control antibody-treated animals. Data are presented as mean ± SEM.

**Table S1.**
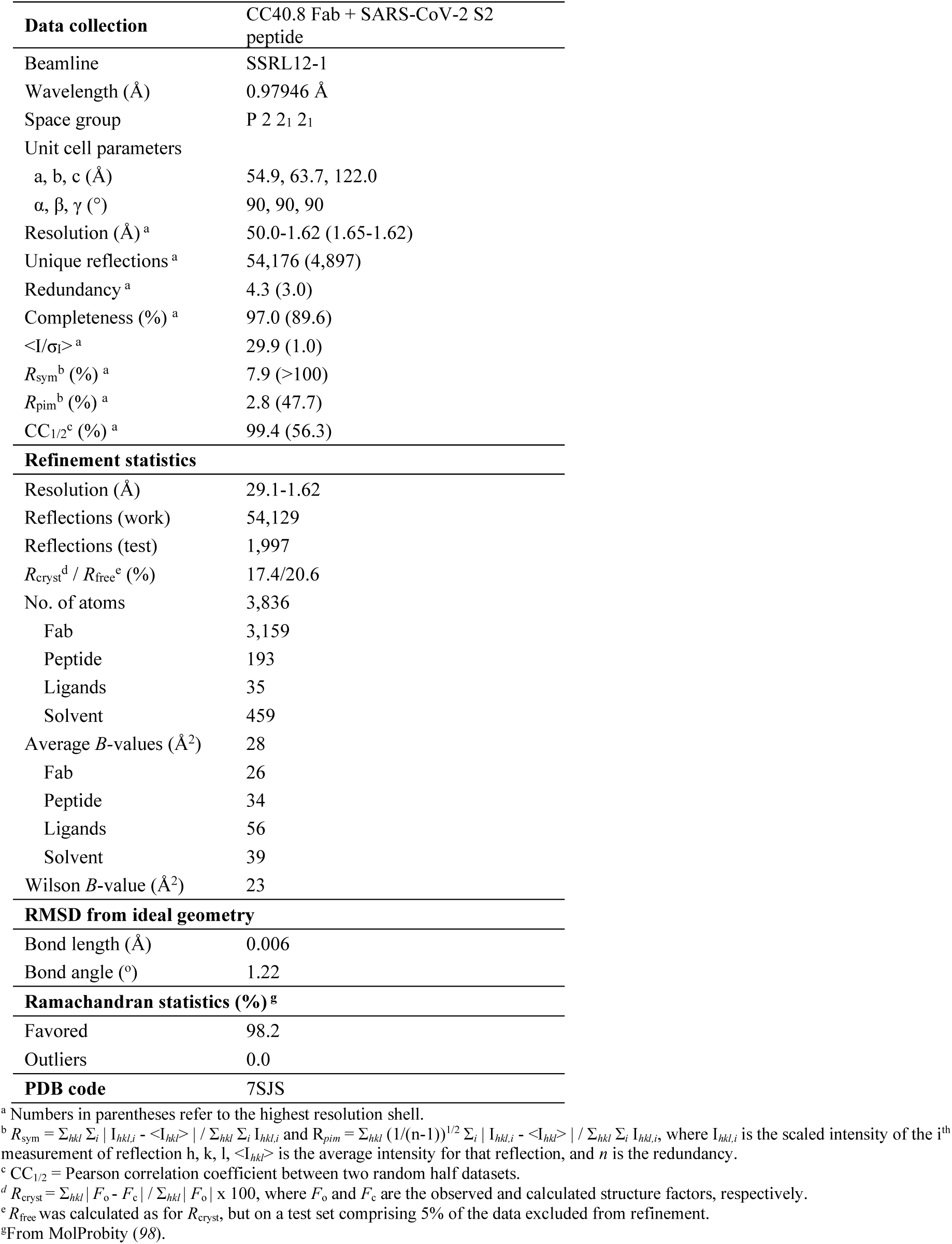
X-ray data collection and refinement statistics.

**Table S2.** Demographic information of COVID-19 convalescent donors.

|  | **COVID donor**<br>**(n = 60)** |
| --- | --- |
| **Age (years)** | 20 to 72 (median = 46) |
| **Gender** |  |
| Male | 47% (28/60) |
| Female | 53% (32/60) |
| **Race/Ethnicity** |  |
| White, non-Hispanic | 80% (48/60) |
| Hispanic | 8.3% (5/60) |
| Black, non-Hispanic | 1.7% (1/60) |
| Asian, non-Hispanic | 3.3% (2/60) |
| Unknown | 6.7% (4/60) |
| SARS-CoV-2 PCR Positivity | 75% (45/60) |
| Lateral Flow Positivity | 60% (36/60) |
| **Disease Severity** |  |
| Mild | 56.7% (34/60) |
| Mild to Moderate | 6.7% (4/60) |
| Moderate | 25% (15/60) |
| Moderate to Severe | 5% (3/60) |
| Severe | 5% (3/60) |
| Critical | 1.7% (1/60) |
| **Symptoms** |  |
| Cough | 60% (36/60) |
| Fever | 55% (33/60) |
| Fatigue | 38.3% (23/60) |
| Anosmia | 31.7% (19/60) |
| Dyspnea | 26.7% (16/60) |
| Diarrhea | 16.7% (10/60) |
| Days Post Symptom Onset<br>at Collection | 6 to 90 (median = 35.5) |

